# Targeting Glioblastoma Signaling and Metabolism with A Re-Purposed Brain-Penetrant Drug

**DOI:** 10.1101/2021.03.16.435487

**Authors:** Junfeng Bi, Atif Khan, Jun Tang, Sihan Wu, Wei Zhang, Ryan C. Gimple, Tomoyuki Koga, Aaron M. Armando, Shunichiro Miki, Huijun Yang, Briana Prager, Ellis J. Curtis, Derek A. Wainwright, Frank B. Furnari, Jeremy N. Rich, Timothy F. Cloughesy, Oswald Quehenberger, Harley I. Kornblum, Andrey Rzhetsky, Benjamin F. Cravatt, Paul S. Mischel

## Abstract

The highly lethal brain cancer glioblastoma (GBM) poses a daunting challenge because the blood-brain barrier renders potentially druggable amplified or mutated oncoproteins relatively inaccessible. Here, we identify SMPD1, an enzyme that regulates the conversion of sphingomyelin to ceramide and a critical regulator of plasma membrane structure and organization, as an actionable drug target in glioblastoma. We show that the safe and highly brain-penetrant antidepressant fluoxetine, potently inhibits SMPD1 activity, killing GBMs, in vitro and in patient-derived xenografts, through inhibition of EGFR signaling and via activation of lysosomal stress. Combining fluoxetine with the chemotherapeutic agent temozolomide, a standard of care for GBM patients, causes massive increases in GBM cell death, and complete and long-lived tumor regression in mice. Incorporation of real-world evidence from electronic medical records from insurance databases, reveals significantly increased survival in glioblastoma patients treated with fluoxetine, which was not seen in patients treated with other SSRI anti-depressants. These results nominate the repurposing of fluoxetine as a potentially safe and promising therapy for GBM patients and suggest prospective randomized clinical trials.

## Introduction

The highly lethal brain tumor glioblastoma (GBM) is one of the most difficult forms of cancer to treat. Despite a relatively advanced catalogue of the mutational landscape of GBM, genomic insights have failed to translate into improved survival for the vast majority of patients, most of whom still die within two years, despite aggressive treatment with surgical resection, radiotherapy, and temozolomide (Cloughesy et al., 2014). Multiple challenges contribute to persistent therapeutic failure. First, many targeted cancer drugs have relatively poor brain/plasma ratios, resulting in systemic toxicities that preclude adequate target inhibition in patients. Second, the underlying biology of actionable genetic alterations in the brain appears to be profoundly influenced by the brain’s unique physiology in ways that are not well understood (Bi et al., 2020; Brennan et al., 2013; Mack et al., 2016; Nagaraja et al., 2019; Quail and Joyce, 2017). Third, glioblastomas commonly contain extrachromosomal DNA (ecDNA), in which growth-promoting oncogenes, including the epidermal growth factor receptor (EGFR), are amplified at very high levels (Kim et al., 2020; Morton et al., 2019; Nathanson et al., 2014; Nikolaev et al., 2014; Turner et al., 2017; Xu et al., 2019; Zhou et al., 2017). Unfortunately, reversible and rapid modulation of the level of these ecDNAs appears to play a key role in driving GBM resistance to targeted therapy (Nathanson et al., 2014), thereby motivating a search for alternative treatment strategies.

The palette of genomic alterations in GBM appears to differ in some consistent ways from that observed in other cancers that don’t arise in the brain. For example, EGFR kinase domain mutations, which more commonly occur in other types of systemic cancers, are very rare in GBMs (Brennan et al., 2013; Sanchez-Vega et al., 2018), in which instead *EGFR* amplification is a dominant oncogenic mechanism. Under normal physiological conditions, EGFR ligands promote dimerization and downstream signaling (Arkhipov et al., 2013; Lemmon et al., 2014). In GBM, the amplified *EGFR*s on ecDNA often contain mutations in the extracellular domain of the receptor, such as EGFRvIII, which disrupt ligand binding, but nonetheless promote oncogenic signaling, raising the possibility that something unique about the brain’s microenvironment may select for *EGFR* amplifications. This motivated us to consider whether the altered lipid environment in the brain, and potentially in tumor cells, might generate a unique selection pressure in GBMs that may expose actionable vulnerabilities. We focused on lipids because recent work has shown that GBM cells may have profoundly altered compositions of cholesterol and phospholipids in the plasma membrane that may determine how EGFRs signal in tumor cells (Bi et al., 2020; Bi et al., 2019; Guo et al., 2011; Villa et al., 2016). We were particularly motivated to search for alterations in sphingolipid biosynthesis pathways, because the balance between sphingomyelin and ceramide is thought to be critical for plasma membrane organization, including the clustering of signaling molecules into discrete membrane domains called lipid rafts (Hannun and Obeid, 2018; Lingwood and Simons, 2010; Ogretmen, 2018; van Meer et al., 2008). We also searched for highly brain-penetrant drugs that selectively and effectively target key enzymatic components of the sphingolipid biosynthesis machinery.

Here, we proceed from unbiased identification and validation of acid sphingomyelinase (SMPD1) as a compelling GBM target that is required for tumor cell survival, to dissection of its underlying actionable enzymatic mechanism, to identification of fluoxetine as an FDA-approved, safe and highly brain-penetrant drug that potently inhibits SMPD1 for the treatment of patients with glioblastoma. We conduct *in vivo* proof of concept studies revealing complete tumor regression in patient-derived GBM brain tumor models in mice, when fluoxetine is combined with standard of care. Fluoxetine (Prozac) has been prescribed for years (Wong et al., 2005). Therefore, we realized that there may be an opportunity to glean real-world evidence for the potential therapeutic benefit of fluoxetine, which have been shown to be highly brain-penetrant and safe. We complement our experimental data with analyses from electronic medical records demonstrating that combining fluoxetine specifically with standard of care, unlike other SSRIs analyzed, significantly prolongs survival in patients with glioblastoma.

## Results

### Glioblastomas Highly Depend on SMPD1 for Survival

Hypothesizing that sphingolipid metabolism may play an important role in glioma pathogenesis (Bi et al., 2020; Noack et al., 2014; Ogretmen, 2018), we analyzed a large-scale RNA interference cancer dependency dataset (DepMap) (DepMap, 2020; McFarland et al., 2018) containing over 600 cancer cell lines of different histological types, including 43 glioma cell lines. We focused on 14 genes that encode the key enzymes in the sphingolipid synthesis pathway (Figures 1A and S1A) and are coordinately upregulated in glioblastoma clinical samples (Figures S1B and S1C). We identified sphingomyelin phosphodiesterase 1 (SMPD1), also known as acid sphingomyelinase (ASM), as the top survival dependency among these 14 genes in glioma cell lines (Figures 1B, S1D, and S1E). Concordant with its potential role in driving tumor growth, elevated *SMPD*1 expression is associated with significantly shorter survival in GBM patients from multiple public cancer patient datasets (Figures 1C and S1F-S1H).

**Figure 1.**
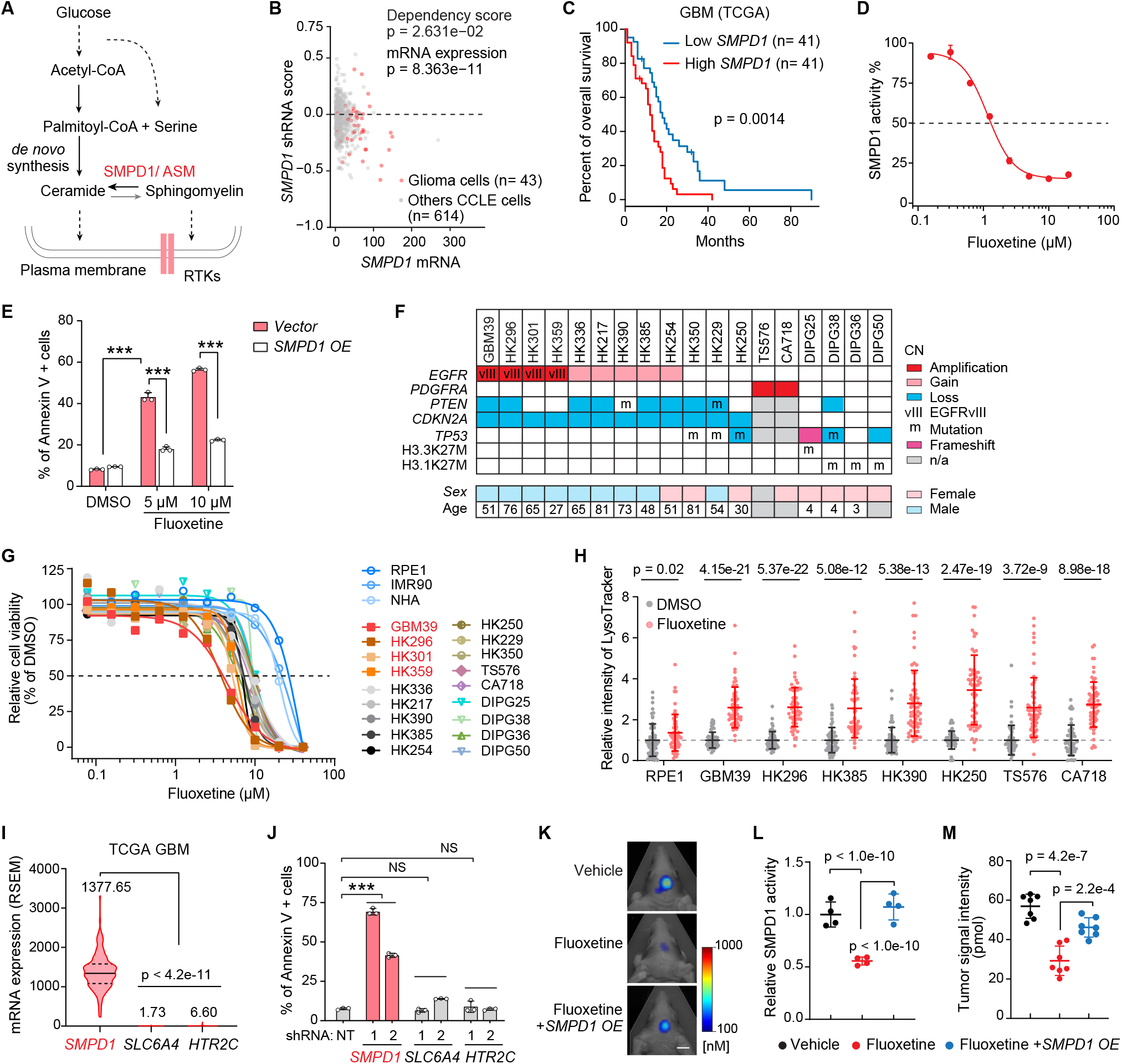
GBMs Depend on SMPD1 for Survival, Making Them Sensitive to Fluoxetine-Mediated Cell Death. (A) Schematic pathway of sphingolipid metabolism in plasma membrane lipid remodeling of GBM cells. (B) shRNA effect scores and mRNA levels of *SMPD1* in glioma and other CCLE cell lines from the DepMap dataset. (C) Kaplan-Meier analysis of overall survival of GBM patients with high or low *SMPD1* mRNA expression in the TCGA GBM (RNA-seq) dataset. (D) Enzymatic activity of SMPD1 in U87EGFRvIII cells with 24 hours fluoxetine treatment. (E) Percentage of Annexin V-positive cells in DMSO or fluoxetine treated U87EGFRvIII cells. (F) Brief information, including major genomic features and sex, of 18 patient-derived GBM neurosphere lines. CN, copy number; n/a, not available. (G) Cell viability curves of 3 non-cancer cell lines (NHA, RPE1, and IMR90) and 18 GBM neurosphere lines in response to fluoxetine treatment (n= 4). (H) LysoTracker staining in indicated cell lines (n= 60). (I) mRNA level (RSEM) in TCGA GBM patient samples. (J) Percentage of Annexin V-positive U87EGFRvIII cells with indicated shRNA knockdown (NT, non-targeting). (K-M) Representative tumor images (K), SMPD1 enzymatic activity (n= 4) (L), and tumor signal intensity (n= 7) (M) of U87EGFRvIII orthotopic xenograft models. Scale bar, 5 mm. Data represent mean± SD except (I). The median value (center line) and the 25th and 75th percentiles (dash lines) are presented in (I). Two-tailed Student’s t-test for (B) and (H). Log-rank test for (C). ANOVA followed by Tukey’s multiple comparisons test for (E), (I), (J), (L), and (M). ***p < 0.001; NS, not significant.

### Pharmacological Inhibition of SMPD1 by Fluoxetine Selectively Kills GBMs

SMPD1 catalyzes the conversion of sphingomyelin to ceramide (Hannun and Obeid, 2018). Complete genetic loss of SMPD1, which occurs in children with Niemann Pick disease (Schuchman and Desnick, 2017), results in elevated sphingomyelin levels, lysosomal stress, and cell death in some contexts (Hannun and Obeid, 2018; Schuchman and Desnick, 2017). To determine whether GBM cells, because of their enhanced dependence on SMPD1, could potentially be highly sensitive to a pharmacological inhibitor of SMPD1, we searched the literature for FDA-approved, brain penetrant drugs that have been shown to inhibit SMPD1 enzymatic activity. The selective serotonin reuptake inhibitor (SSRI) anti-depressant fluoxetine (Prozac) was recently identified as a potential SMPD1 inhibitor (Gulbins et al., 2013; Kornhuber et al., 2008). In glioblastoma cells, fluoxetine inhibited SMPD1 enzymatic activity (Figure 1D), resulting in dose-dependent glioblastoma cell death (Figures 1E and S2A-S2D). *SMPD1* overexpression abrogated the effect of fluoxetine on glioblastoma cells, supporting on-target activity (Figures 1E, S2E, and S2F). To further determine the anti-GBM potential of fluoxetine, we performed a sensitivity screen in 3 non-cancer cell lines and 18 patient-derived glioblastoma cultures of various tumor genotypes (Figure 1F and Table S1). Fluoxetine resulted in tumor-specific cell death (Figure 1G), and caused extensive lysosomal stress in the patient-derived GBM cultures (Figure 1H), as would be predicted as a marker for an SMPD1 inhibitor (Schuchman, 2010).

To determine whether the anti-tumor effect could be mediated through the serotonin reuptake system, we analyzed the serotonin transporter SLC6A4 and the serotonin receptor HTR2C. Neither gene was expressed at appreciable levels in glioblastoma clinical samples (Figure 1I), and shRNA knockdown of either gene did not significantly affect glioblastoma viability, in contrast to shRNA knockdown of *SMPD1*, which caused substantial glioblastoma cell death (Figures 1J and S2G). Other serotonin receptors also failed to show appreciable transcript levels or survival association in the TCGA GBM dataset (Figures S2H and S2I). Importantly, fluoxetine administration significantly inhibited tumor growth, in accordance with dramatic inhibition of SMPD1 enzymatic activity, in orthotopic xenografts implanted in the brain of nude mice (Figures 1K-1M), which were rescued by *SMPD1* overexpression (Figures 1K-1M). These data do not exclude a potential modulatory role for serotonergic activity in the tumor microenvironment (Caudill et al., 2011; Dolma et al., 2016; Mahe et al., 2004), but they do further suggest that fluoxetine kills GBM cells through an alternative mechanism, including SMPD1 inhibition *in vitro* and *in vivo*. Together, these results suggest that GBMs depend on SMPD1 for survival and are highly sensitive to fluoxetine-mediated SMPD1 inhibition.

### Fluoxetine Kills GBM Cells by Disrupting Sphingomyelin Metabolism with Resultant Inhibition of Oncogenic EGFR Signaling

Many GBMs contain amplified *EGFRvIII*, a constitutively active form mutation of *EGFR* driving GBM malignant progression (Cloughesy et al., 2014). In our sensitivity screen, we found that GBMs with *EGFRvIII* amplification were significantly more sensitive to fluoxetine than other GBMs (Figure 2A). Further, overexpression of EGFRvIII in a GBM cell line globally increased sphingolipids levels (Figures S2J and S2K), sensitized tumor cells to an inhibitor of sphingolipid *de novo* synthesis (Figures S2L-S2N) as well as *SMPD1* shRNA knockdown (Figures S2O-S2Q), and generated dose-dependent sensitivity to fluoxetine (Figures S2A-S2C). Patient-derived GBMs with endogenously amplified EGFRvIII were similarly, highly sensitive to *SMPD1* depletion or fluoxetine, both in neurosphere cultures (Figures 2B-2D and S2R) and in orthotopic xenografts implanted in the brain of nude mice (Figures 1K-1M), further suggesting a potential role of EGFRvIII or downstream oncogenic signaling in the anti-GBM effect of fluoxetine.

**Figure 2.**
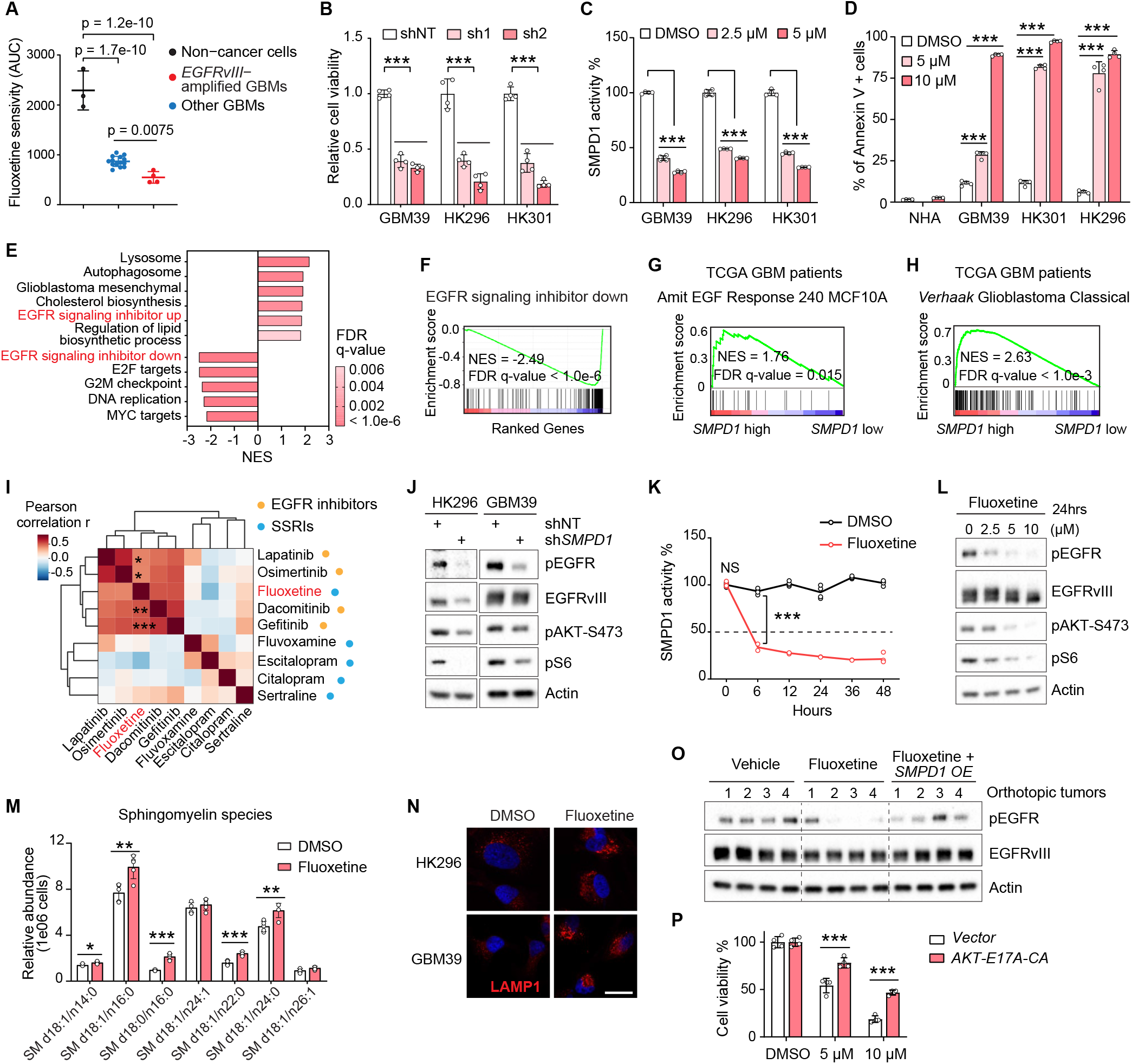
Fluoxetine’s Inhibition of SMPD1 Blocks Oncogenic EGFR Signaling in GBM Cells. (A) Fluoxetine sensitivity (area under the cell viability curve) of 3 non-cancer cell lines and 18 patient-derived GBM neurospheres, including 4 *EGFRvIII*-amplified lines. (B) Relative cell viability of 3 *EGFRvIII*-amplified GBM neurosphere lines with *SMPD1* or non-targeting shRNAs (n= 4). (C) SMPD1 enzymatic activity in GBM neurosphere lines with 24 hours of fluoxetine treatment (n= 4). (D) Flow cytometry analysis of Annexin V-positive cells in normal human astrocytes (NHA) and 3 GBM neurosphere lines (n= 4). (E and F) Gene Set Enrichment Analysis identifies differentially enriched or depleted transcripts in 3 glioblastoma neurosphere cultures treated with fluoxetine vs DMSO. (G and H) Gene Set Enrichment Analysis of differentially expressed genes in GBM clinical samples (TCGA, HUG133A GBM) with high or low *SMPD1* expression. (I) Drug sensitivity correlation of fluoxetine, 4 EGFR inhibitors, and 4 other SSRI antidepressants in all 40 glioma cell lines from the DepMap dataset. (J) Western blot analysis of EGFR signaling in HK296 and GBM39 cells. (K) Relative enzymatic activity of SMPD1 in GBM39 cells treated with DMSO or 5 µM fluoxetine for indicated hours (n= 4). (L) Western blot analysis of EGFR signaling activity in GBM39 cells with indicated treatments for 24 hours. (M) Lipidomic analysis of sphingomyelin species in U87 cells expressing EGFRvIII with 5 µM fluoxetine or DMSO treatment (n= 5). (N) LAMP1 staining of GBM cells with DMSO or fluoxetine treatment. Scale bar, 20 µm. (O) Western blot analysis of EGFR phosphorylation in orthotopic GBM xenograft tumors. (P) Viability of GBM39 cells expressed vector or a constitutively active AKT E17A-CA allele (n= 4). Data represent Mean± SD. Two-tailed Pearson for (I). Two-tailed Student’s t-test for (M). ANOVA followed by Tukey’s multiple comparisons test for (A-D), (K), and (P). *p < 0.05; **p < 0.01; ***p < 0.001; NS, not significant.

RNA sequencing in these three *EGFRvIII* amplified, patient-derived GBM neurosphere lines treated with fluoxetine for 42 hours, revealed a transcriptional signature indicative of EGFR inhibition (Figures 2E, 2F, and S3A-S3D). Analysis of TCGA GBM clinical samples revealed a highly significant correlation between *EGFR* amplification, the transcriptional signature of EGFR signaling, and *SMPD1* expression (Figures 2G, 2H, and S3E), suggesting a potential molecular basis for enhanced fluoxetine sensitivity that was consistent with enhanced SMPD1 survival dependency in glioma cell lines with elevated EGFR protein levels. Interestingly, in the merged cohort of low-grade glioma and GBM patients, deep deletion of *SMPD1* and *EGFR* amplification/mutations are mutually exclusive (p < 0.001) (Figure S3F), suggesting a synthetic lethal interaction between *SMPD1* and *EGFR*. We further noted that, surprisingly, in an analysis of drug sensitivity dataset from DepMap among all 40 glioma cell lines, fluoxetine clustered with 4 bona fide EGFR inhibitors, not with 4 other SSRI anti-depressants (Figure 2I), further indicating the anti-GBM activity of fluoxetine may be through inhibiting EGFR signaling, but not serotonin transporters.

Next, we set out to determine how fluoxetine might affect EGFR signaling. First, we confirmed the effect of *SMPD1* genetic depletion on suppressing EGFRvIII signaling (Figures 2J and S4A). Fluoxetine rapidly inhibited SMPD1 enzymatic activity (Figure 2K), but inhibition of EGFRvIII phosphorylation and downstream signaling became apparent only after approximately 24 hours (Figures 2L, S4B, and S4C). This timing suggests an indirect and delayed, but potentially important mechanism of EGFR inhibition. We found a significant increase in sphingomyelin levels, with SM d18:1/n16:0 and SM d18:1/n24:0 being the most abundant species, in response to fluoxetine induced SMPD1 inhibition (Figures 2M and S4D-S4F). Lysosomal stress, measured by LAMP1 and lysotracker staining (Figures 2N, S4G, and S4H), was also detected. Further, overexpression of *SMPD1* rescues the EGFR signaling inhibited by fluoxetine both in GBM cell lines and in orthotopic GBM tumors (Figures 2O and S4I). To determine whether the loss of EGFRvIII signaling contributed to fluoxetine’s anti-GBM activity, we focused on AKT, which has been shown as a major signaling output that is required for EGFRvIII’s oncogenic effects(Cloughesy et al., 2014). Expression of the constitutively active AKT E17A-CA allele significantly rescued the anti-tumor effect of fluoxetine (Figures 2P, S4J, and S4K). Taken together, these results confirm that fluoxetine selectively kills GMB cells by disrupting SMPD1-mediated sphingomyelin metabolism and inhibiting oncogenic EGFR signaling.

### Disrupting Sphingomyelin Metabolism Inhibits EGFR Activity on Plasma Membrane of GBM Cells

SMPD1 inhibition, by altering sphingomyelin levels, could potentially affect the structural organization of the plasma membrane, including the highly ordered microdomains, referred to as lipid rafts, in which much signal transduction is through to occur (Arkhipov et al., 2013; Bi et al., 2019; Sezgin et al., 2017). Therefore, we analyzed the effect of fluoxetine on membrane order in live GBM cells by using the lipid phase-sensitive fluorescent probe Laurdan (Owen et al., 2011; Parasassi et al., 1997). Laurdan staining quantifies shifts in the emission spectra generated by probe binding to ordered vs disordered phases in the plasma membrane (Owen et al., 2011). Fluoxetine treatment lowered the generalized polarization (GP) values of the plasma membrane (Figures 3A and 3B). *SMPD1* overexpression reversed the effect of fluoxetine on membrane order, thereby demonstrating a direct effect on tumor cell plasma membrane architecture (Figures 3A and 3B). In line with the recent finding that SMPD1 inhibition caused KRAS mislocalization from the plasma membrane (Cho et al., 2016; Schuchman, 2010), fluoxetine treatment resulted in depletion of EGFRvIII from the plasma membrane with loss of downstream EGFRvIII signaling (Figures 3C and 3D), which was rescued by *SMPD1* overexpression, both *in vitro* and *in vivo* (Figures 2O and S4I). Concordant with these changes, we observed a dramatic loss of EGFRvIII from the raft marker-enriched membrane fraction (Figures 3E and 3F) in fluoxetine-treated GBM cells, as well as an enhanced EGFR internalization from the plasma membrane in fluoxetine-treated GBM cells (Figure 3G). This was followed by reduced EGFRvIII protein levels, which was partially rescued by proteasome and lysosome inhibitors (Figures S5A-S5D), potentially explaining the reduced levels of EGFRvIII protein, as well as the reduced levels of pEGFRvIII and its downstream effectors (Figures 2L, S4B, and S4C).

**Figure 3.**
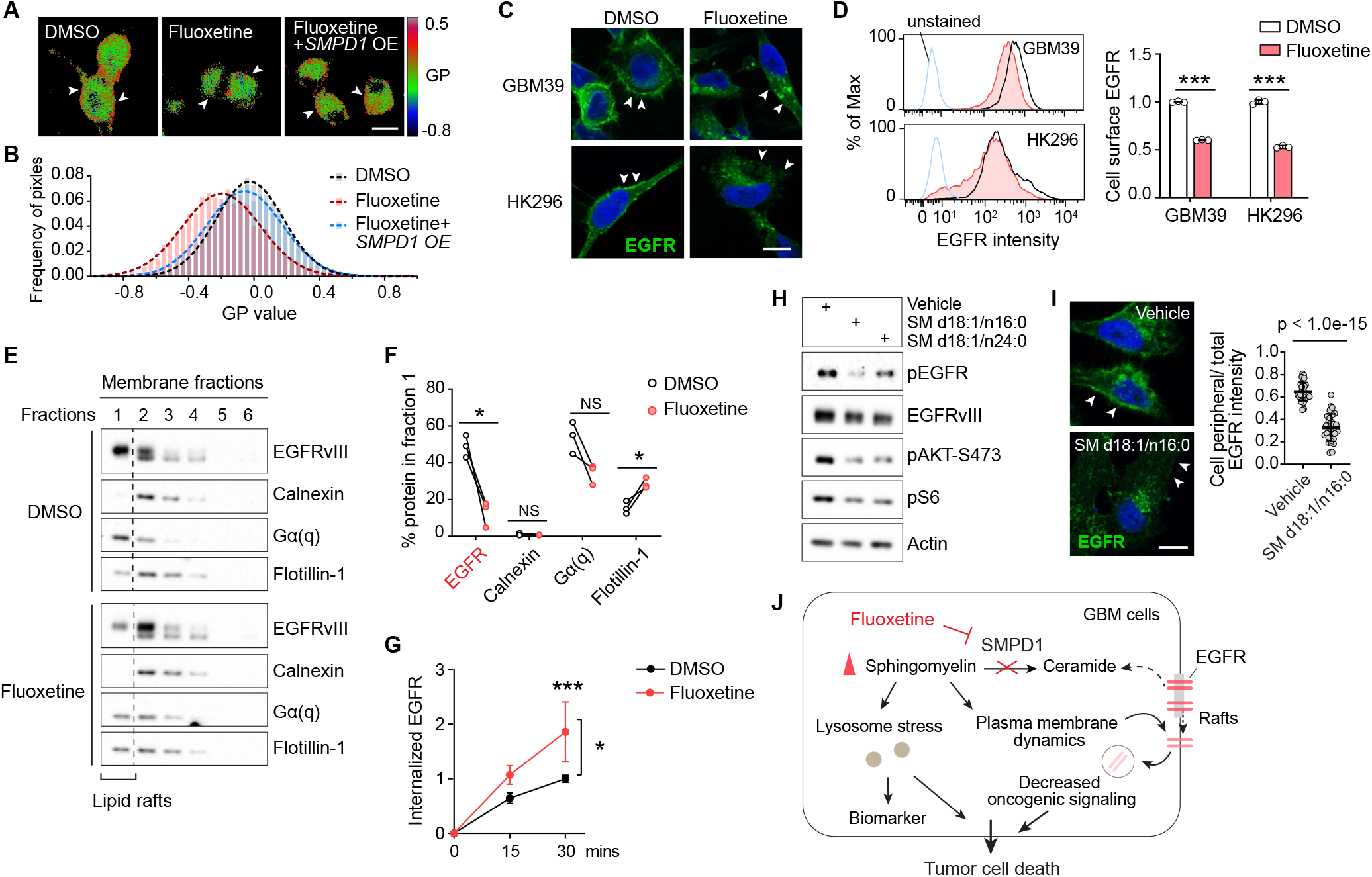
By Increasing Sphingomyelin Levels, Fluoxetine Causes Loss of Cell Surface EGFR from Membrane Rafts with Subsequent Receptor Internalization and Degradation. (A and B) Laurdan imaging analysis of membrane lipid order in U87 cells expressing EGFRvIII at baseline, after fluoxetine treatment, and with overexpression of an *SMPD1* construct. Generalized polarization (GP) images indicate higher membrane order (red) and lower membrane order (blue). Scale bar, 20 µm. (C and D) Imaging and flow cytometry analysis (n= 3) of cell surface EGFR in GBM cells. Scale bar, 10 µm. (E) Western blot analysis of EGFRvIII and marker proteins in the membrane fractions of GBM39 cells. Calnexin is a marker for non-lipid rafts fractions, while Gα(q) and Flotillin-1 are markers of lipid rafts. (F) Percentage of indicated protein levels in fraction 1, the lipid rafts fraction, which is absent with non-lipid rafts marker Calnexin and present with lipid raft markers Gα(q) and Flotillin-1. Data were normalized to total protein levels of all six fractions (n= 3). (G) Internalized EGFR of GBM39 cells by flow cytometry (n= 4). (H) EGFR signaling in GBM39 cells treated with sphingomyelins or vehicle. (I) EGFR antibody staining showing loss of cell surface EGFR in GBM39 cells after SM d18:1/n16:0 treatment. Scale bar, 10 µm. (J) Schematic model of the fluoxetine-SMPD1 axis in regulating sphingomyelin metabolism and oncogenic receptor signaling of GBM cells. Data represent Mean± SD. Two-tailed Student’s t-test for (D), (F), and (I). ANOVA followed by Tukey’s multiple comparisons test for (G). *p < 0.05, ***p < 0.001, NS, not significant.

To further confirm that fluoxetine’s effect on EGFRvIII signaling was mediated by SMPD1 inhibition, we added SM d18:1/n16:0 and SM d18:1/n24:0, the two sphingomyelins that were dramatically elevated after fluoxetine treatment (Figures 2M and S4D-S4F) to GBM cells. Both sphingomyelins significantly suppressed EGFRvIII signaling and GBM cell viability, which was further enhanced by fluoxetine (Figures 3H, and S5E-S5H). Ceramide was not able to rescue these effects (Figure S5E). Further, the addition of SM d18:1/n16:0 caused loss of EGFRvIII from the plasma membrane of GBM cells (Figure 3I), mimicking the effects of fluoxetine. Taken together, these results suggest that fluoxetine causes loss of EGFR from the plasma membrane and inhibits EGFR signaling by blocking SMPD1 enzymatic activity and elevating sphingomyelin levels (Figure 3J).

### Fluoxetine Efficacy in Patient-Derived Orthotopic GBM Mice Models

Fluoxetine is highly brain-penetrant (Bolo et al., 2000; Karson et al., 1993) and is FDA approved for a variety of neuropsychiatric disorders (Eli Lilly and Company, 2017; Wong et al., 2005). It has been demonstrated to be safe over a range of doses from 20-80 mg/day, with most depression patients being treated with the lower doses (Eli Lilly and Company, 2017). To better understand how fluoxetine could potentially be used as a treatment, we tested the calculated mouse equivalent oral doses of the FDA-approved dose range, in patient-derived, *EGFRvIII* amplified, glioblastomas implanted into the brains of nude mice (Figures 4A and S6A). In the GBM39 model, we observed significant, dose-dependent tumor growth inhibition and markedly prolonged mouse survival at mouse dose equivalents of 50 and 80 mg/day (Figures 4B, 4C, and S6B). No toxicity was observed (Figure S6C). The low-dose fluoxetine treatment, 4.2 mg/kg, which translates to 20 mg per day in humans, did not affect tumor growth (Figures 4B and 4C). In a second, independent patient-derived, *EGFRvIII* amplified, glioblastoma model implanted into the brains of nude mice, HK296, daily treatment with 10 mg/kg and 15 mg/kg significantly inhibited glioblastoma growth (Figures 4D and 4E), blocked tumor cell proliferation (Figures 4F and S6D), induced tumor cell death (Figure 4G and S6D), and markedly prolonged mouse survival (Figure 4H), concomitant with EGFR inhibition and increased lysosomal stress (Figures 4I-4K, S6E, and S6F). Importantly, fluoxetine-treated mice showed no evidence of toxicity, no weight loss, and no cell death or elevation of LAMP1 in the surrounding brain, using antibodies that detect both human and mouse protein (Figures 4G and S6C-S6F). These results demonstrate that clinically safe and achievable doses of fluoxetine that can inhibit SMPD1 activity may potentially be used to treat glioblastoma patients and that there is likely to be a relatively wide therapeutic window.

**Figure 4.**
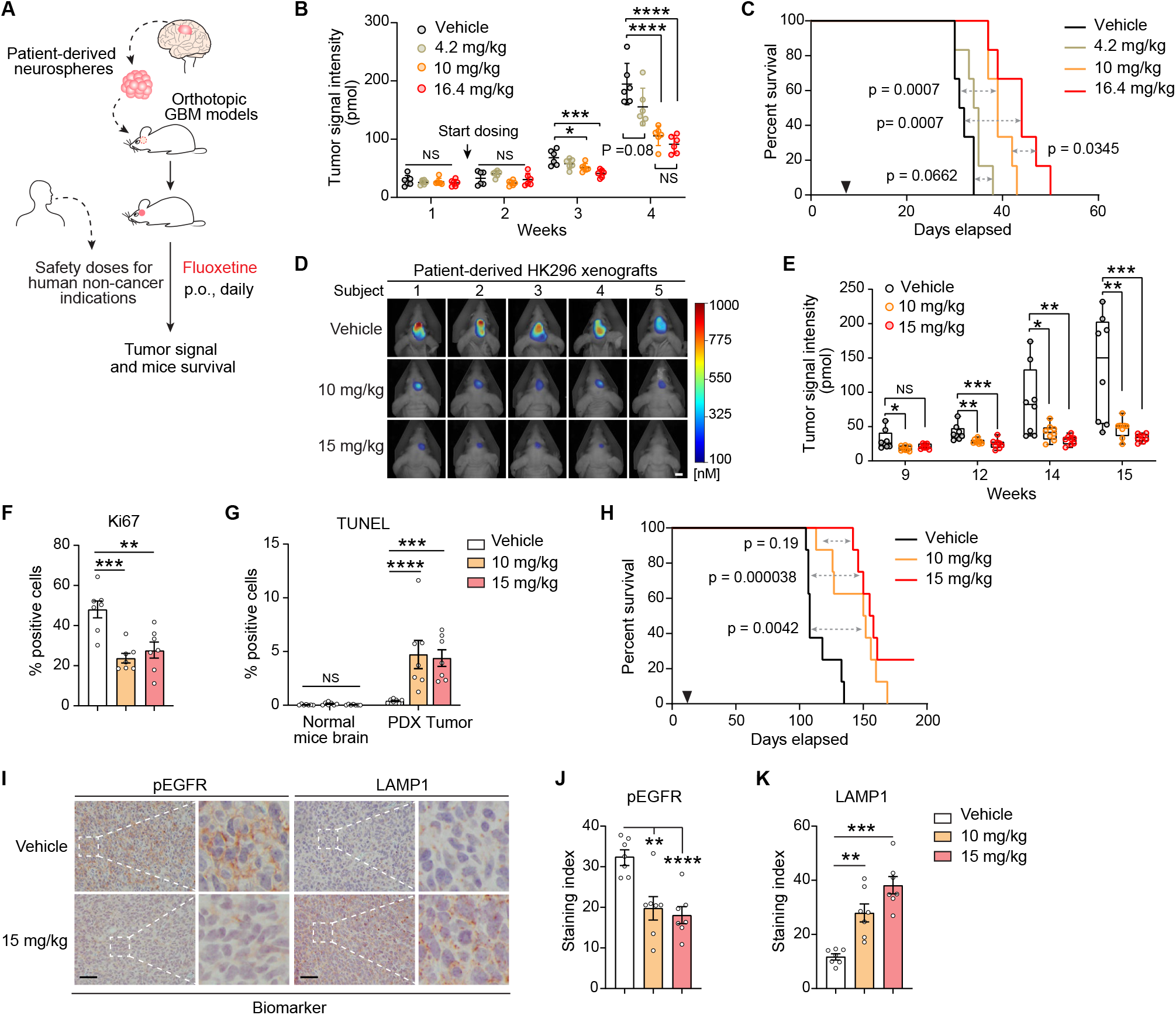
Fluoxetine Promotes Tumor Regression and Prolongs Survival of Mice Bearing Patient-Derived Orthotopic GBMs. (A) Schematic of the fluoxetine treatment protocol in patient-derived GBM orthotopic xenograft mouse models. (B and C) Tumor signal intensity (B) and Kaplan-Meier survival analysis (C) of patient-derived GBM39 orthotopic xenograft models with vehicle or fluoxetine administrations (n= 6, p.o., daily). Safety doses of fluoxetine for human non-cancer indications were converted to mice doses based on body surface area. 4.2 mg/kg in mice is equal to the minimal suggested dose for human indications, and 16.4 mg/kg in mice is equal to the maximal dose for human indications. (D and E) Representative tumor images at week 15 (D) and tumor signal intensity (E) of patient-derived HK296 orthotopic xenograft models with vehicle or fluoxetine administrations (n= 8, p.o., daily). Scale bar, 5 mm. The median value (center line), the min and max (whiskers), and the 25th and 75th percentiles (box perimeters) are presented. (F) Percentage of Ki67 positive cells in HK296 xenograft tumors. (G) Percentage of TUNEL positive cells in HK296 xenograft tumors and surrounding mice brains. (H) Kaplan-Meier survival analysis of mice bearing HK296 xenograft tumors (n= 8). (I-K) Immunohistochemistry analysis of two biomarkers, phosphorylated-EGFR, and LAMP1, in HK296 xenograft tumors. Scale bar, 50 µm. Data represent mean± SD in (B) and ± SEM in (F), (G), (J), and (K). ANOVA followed by Tukey’s multiple comparisons test for (B), (E)-(G), (J), and (K). Log-rank test for (C) and (H). *p < 0.05; **p < 0.01; ***p< 0.001; ****p < 0.0001; NS, not significant.

### Combining Fluoxetine with Temozolomide Suppresses GBM Recurrence and Prolongs Survival

Currently, most glioblastoma patients, including those with EGFRvIII amplification, are treated with the alkylating chemotherapy temozolomide (TMZ), in addition to surgery and radiotherapy. Although the effects of fluoxetine monotherapy were clear, they were not overwhelming, and we hypothesized that fluoxetine might synergize temozolomide because of the potential role for EGFR signaling in regulating DNA damage repair (Squatrito and Holland, 2011). We also found that fluoxetine treatment resulted in the downregulation of genes in DNA repair pathway in GBM cells (Figures S7A and S7B). *In vitro*, fluoxetine was highly synergistic with temozolomide in inducing DNA damage and cell death (Figures 5A, 5B, and S7C-S7F). This combination effect was mediated through downstream EGFRvIII signaling because the AKT E17A-CA allele rescued the cell death (Figure S7F).

**Figure 5.**
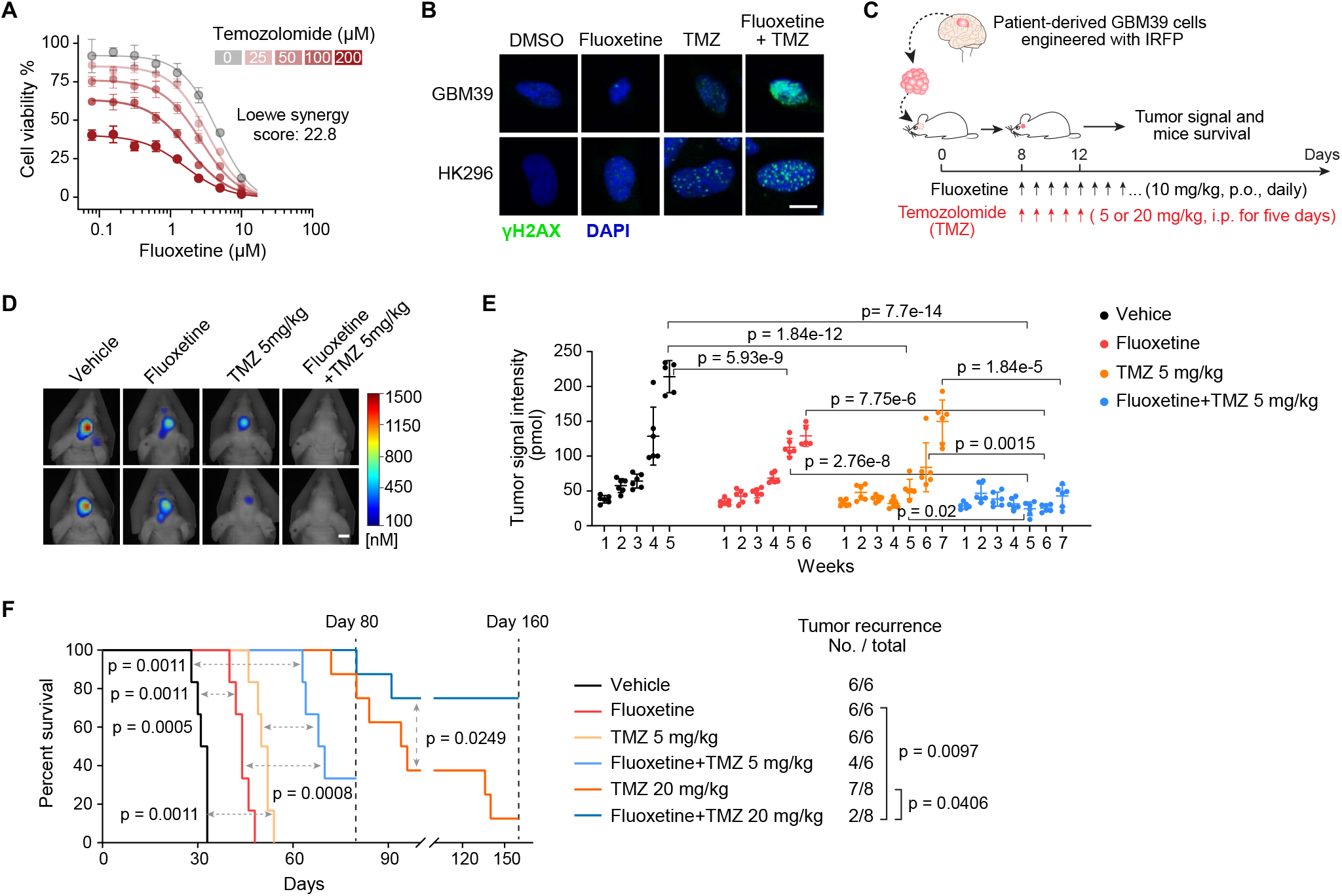
Combining Fluoxetine with Temozolomide Suppresses GBM Recurrence and Prolongs Survival. (A) Synergistic effect of fluoxetine and temozolomide (TMZ) in U87 cells with EGFRvIII overexpression. Data represent mean± SD of four biological replicates. (B) γH2AX staining of GBM39 and HK296 cells with indicated treatments. 2.5 µM for fluoxetine, and 50 µM for TMZ. Scale bar, 10 µm. (C) Schematic overview of the fluoxetine-TMZ combination therapy in patient-derived GBM39 orthotopic xenograft models. (D-F) Representative tumor images at week 5 (D), tumor signal intensity (E), and Kaplan-Meier survival analysis (F) of patient-derived GBM39 orthotopic xenograft models with indicated administrations (n= 6 or 8 mice per group). Scale bar, 5 mm. Data represent mean± SD. ANOVA followed by Tukey’s multiple comparisons test in (E). Log-rank test for survival and Fisher’s exact test for tumor recurrence in (F).

Therefore, we compared daily oral fluoxetine treatment alone or in combination with temozolomide treatment for 5 days, followed by maintenance on just fluoxetine (Figure 5C). In the GBM39 orthotopic model, daily fluoxetine was as effective as a 5-day course of daily temozolomide at inhibiting tumor growth and prolonging mouse survival (Figures 5D-5F), and the impact of combination therapy was marked. Continuous daily oral 10 mg/kg fluoxetine treatment, in addition to a five-day course of temozolomide, resulted in prolonged suppression of tumor growth (Figures 5C-5E). Most importantly, mouse survival of the group with 5 mg/kg temozolomide combination therapy more than doubled (p = 0.0011), with 2 of 6 mice showing no tumor recurrence at all after 10 weeks of treatment (Figures 5F and S7G). In 20 mg/kg of temozolomide combination therapy group, 6 of 8 mice showed no tumor recurrence at all after 5 months of treatment (Figures 5F and S7H). No evidence of systemic or neural toxicity, no weight loss, and no cell death or elevation of LAMP1 in the surrounding brain was detected. Taken together, these data suggest that adding fluoxetine at a clinically demonstrated safe dose to standard of care temozolomide, followed by fluoxetine maintenance, could have a major effect on tumor progression, recurrence, and survival.

### Combining Fluoxetine with Standard of Care Treatment Improves the Survival of Brain Tumor Patients

Realizing that fluoxetine has been prescribed for years, we wondered whether we could find “Real-world evidence” that exists in the electronic medical records. We started by examining electronic medical records from the IBM MarketScan insurance claims dataset (2003-2017) which documents healthcare encounters of over 180 million American enrollees. The ascertained death status was available for 378,685 enrollees. Because GBM is a rare condition, and because we used very stringent exclusion criteria to identify cases and controls, the sample size of actual analysis was *N* = 238: we choose to sacrifice statistical power over quality (see Supplemental Document S1).

We started by looking for patients who have an ICD9 or 10 (International Classification of Disease code) for “malignant neoplasm of brain”, are over the age of 18, and lack any other cancers that could be metastatic. We then looked only for adult patients who had surgical resection of the tumor along with radiation therapy and temozolomide, to ensure that we are looking at glioblastoma patients (Figure 6A). For a final cohort of 238 GBM patients, we estimated survival probability and hazard ratio of all-cause deaths with and without SSRI exposure after controlling for age, sex, and also for immortal time bias (Levesque et al., 2010; Suissa, 2008). We found that patients who have fluoxetine added to standard of care had significantly longer median overall survival (fluoxetine: 545 days vs control: 318 days). The age and sex adjusted hazard ratio of all-cause death in fluoxetine treated group was 0.42 [95% CI, 0.20 – 0.88], p = 0.02 compared to the control group (Figure 6B). This survival benefit was not found in patients treated with two other SSRIs, citalopram and escitalopram (Figures 6C and 6D), which were also shown not to have activity against glioma cell lines (Figure 2I), further suggesting the anti-GBM activity of fluoxetine is independent of its function as an SSRI and pointing to the unique anti-GBM activity of fluoxetine. Please see Supplemental Document S1 for complete details of these analyses. These results suggest that a combination of fluoxetine with standard of care treatment may help to improve the survival of brain tumor patients (Figure S7I).

**Figure 6.**
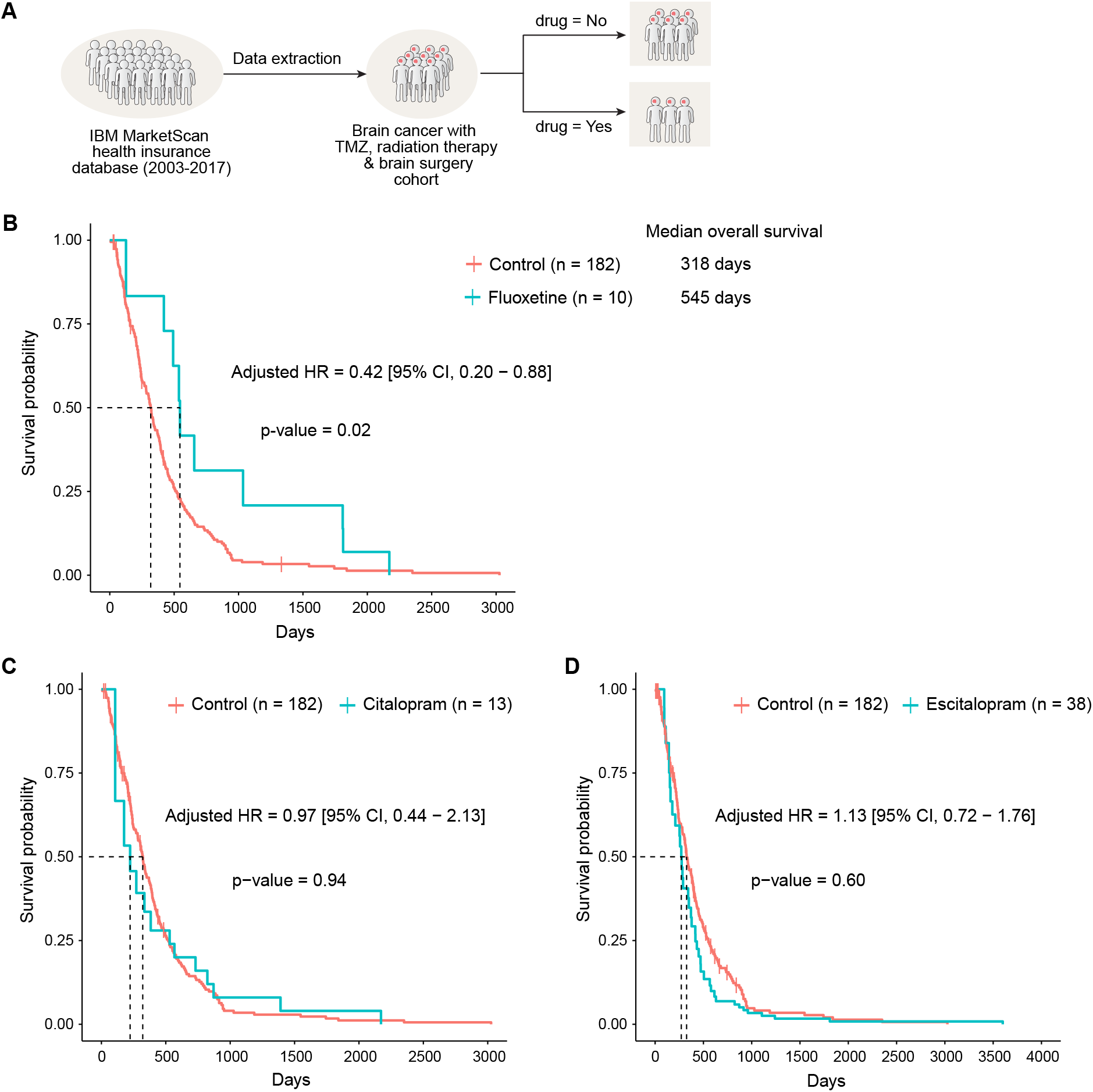
Real-World Electronic Medical Record Evidence for Efficacy and Specificity: Combining Fluoxetine, but Not Citalopram or Escitalopram, Significantly Prolongs Survival of Brain Tumor Patients. (A) Outline of the strategy utilized for define and enrichment of GBM patient cohort in electronic medical records from the IBM MarketScan dataset (2003-2017). (B-D) Survival curve of patients in the GBM-enriched cohort treated with fluoxetine and two other SSRI anti-depressants using time-dependent Kaplan-Meier curves. The adjusted hazards ratio was obtained from the extended Cox proportional hazards model after adjusting for age and sex and using SSRI anti-depressant treatment as a time-dependent variable. Patients treated with fluoxetine (B) generally survive longer (median 545 days) compared to those not treated with any SSRI anti-depressants (median 318 days) and adjusted HR 0.42 [95% CI, 0.20 – 0.88], p=0.02, from extended Cox PH model. There was no significant difference in survival for Citalopram (C) and Escitalopram (D) compared to those not treated with any of the three SSRI anti-depressants considered in this study.

## Discussion

GBM has one of the most well-characterized mutational landscapes of any cancer type. Tumors with amplified, *bona fide*, growth-promoting oncogenes occur in over 50% of GBM patients, presenting extremely compelling drug targets. However, the therapeutic promise of precision oncology has yet to be realized for GBM patients. The challenges include: 1) the poor brain/plasma ratio of many targeted cancer drugs that results in dose-limiting toxicities that preclude effective target inhibition, 2) the unique physiology of the central nervous system that contributes to tumor progression in ways that are only beginning to be understood (Bi et al., 2020; Quail and Joyce, 2017), and 3) the frequency of ecDNA amplification in GBM and the rapid and reversible dynamics it causes, collectively creates significant therapeutic challenges. The hoped for new GBM drugs, FDA approvals and better outcomes for patients, have yet to materialize. Here, by identifying the enhanced dependence of GBMs on SMPD1, showing how it is required for regulating plasma membrane dynamics and oncogenic signaling, and showing that fluoxetine, which has remarkably favorable pharmacokinetic properties and ability effective inhibit SMPD1 enzymatic activity, reveals a unique ability amongst SSRI anti-depressants that can translate into benefit for patients. We have identified a potentially effective new way of treating GBM patients with a safe, repurposed, FDA-approved drug, and determined the molecular mechanistic basis underlying it.

Antidepressants are commonly used in medical practice. So why was this potential therapeutic benefit for GBM patients not detected until now? First, as we have shown here, fluoxetine differs from other SSRI’s with vastly differing effects on patient survival and on inhibiting oncogenic EGFR signaling. Therefore, it is not surprising that the two major studies that have looked at the effect of anti-depressants on the outcome for GBM patients did not find a significant signal; they did not distinguish amongst SSRIs (Caudill et al., 2011; Otto-Meyer et al., 2020). In fact, they would not likely have been able to do so while being sufficiently statistically powered. For a relatively rare cancer such as GBM, no institution alone is likely to have enough patients to perform a sufficiently powered retrospective analysis of the effect of fluoxetine on patient outcome. Cooperative groups are also likely to face similar challenges. That is why we leveraged the power of using real-world evidence from a massive database. We were surprised how few patients with GBM were treated with fluoxetine after diagnosis, which also suggests why it would be so difficult for any institution or even a consortium to have detected this quite strong survival enhancing effect. This is precisely the type of challenge for which gathering observational real-world evidence, as we have done here, is so beneficial and why it may provide therapeutic insights that are poised for rapid clinical testing. In fact, FDA leadership recently suggested that real-world evidence derived from real-world data sources such as electronic health records and insurance databases, may have an important role in complementing, but not replacing, randomized controlled clinical trials (Corrigan-Curay et al., 2018; Jarow et al., 2017; Sherman et al., 2016). We believe that incorporating this type of clinical evidence, greatly strengthens the mechanistic and mouse model data we provide, nominating fluoxetine in combination with standard of care as a treatment for GBM patients and suggesting the need for randomized clinical trials in the near term.

Our paper also provides new and unique insight into the critical dependency of GBMs on SMPD1 and its impact on plasma membrane dynamics, including EGFR signaling. We find a critical link between sphingomyelin metabolism and oncogenic receptor signaling on the plasma membrane of GBM cells. Lipids function as the essential components of the plasma membrane, and their compositions are precisely regulated in forming signal microdomains (Lingwood and Simons, 2010; Sezgin et al., 2017; van Meer et al., 2008). Fluoxetine, by blocking SMPD1, causes sphingomyelin to accumulate, resulting in the loss of EGFR receptors from lipid rafts domains and from the cell surface of tumor cells. Interestingly, inhibiting SMPD1 was reported to similarly cause KRAS loss from the plasma membrane of MDCK cells (Cho et al., 2016). Therefore, it will be important in the future to determine if other amplified growth factor receptors similarly induce SMPD1 dependency in GBM and other cancers. Further, sphingomyelins show interaction with cholesterol in the plasma membrane in regulating membrane properties and many intracellular signaling processes (Das et al., 2014; Endapally et al., 2019; Lingwood and Simons, 2010). SMPD1-mediated sphingomyelin metabolism alters the accessible cholesterol pool of the plasma membrane (Das et al., 2014; Endapally et al., 2019), indicating a potential role of membrane cholesterol in the fluoxetine-SMPD1-EGFR axis. This result is consistent with our previous findings that amplified growth factor receptors in GBM may generate a metabolic dependency on cholesterol and saturated phospholipids (Bi et al., 2020; Bi et al., 2019; Villa et al., 2016).

Fluoxetine was initially developed and approved as a selective serotonin reuptake inhibitor (SSRI) for patients with depression (Wong et al., 2005), and it has been recently shown to inhibit SMPD1 enzymatic activity in some contexts (Gulbins et al., 2013; Kornhuber et al., 2008). Our data in GBM tumor cells indicate that fluoxetine blocks over 60% of SMPD1 enzymatic activity within 6 hours and may act as an indirect inhibitor of EGFR by blocking SMPD1 and elevating sphingomyelin levels. Complete germline loss of *SMPD1* causes lysosomal storage disorders because of excess sphingomyelins on the lysosomal membrane (Schuchman and Desnick, 2017). Lysosomal stress that was detected in GBM tumor cells treated with fluoxetine may also contribute to tumor cell death (Petersen et al., 2013) and may be a main mechanism of sensitization in non-EGFR-driven GBMs. Indeed, we found that GBM cells of many different mutational backgrounds were highly sensitive to fluoxetine, although not quite as sensitive as tumor cells containing amplified EGFRvIII, or wild type EGFR. We hypothesize that this sensitivity is mediated through lysosomal stress, which merits further study. Further, besides the DNA repair pathway, other downstream pathways may also potentially contribute to the sensitivity of GBM cells to fluoxetine-TMZ combination therapy (Ma et al., 2016).

In summary, our findings reveal a potentially actionable mechanism that is poised for therapeutic exploitation, using highly brain-penetrant, FDA-approved drugs. In addition to the experimental studies in GBM neurosphere lines and orthotopic PDX mice models, the real-world evidence provided a unique opportunity to ask whether there is a reason to think that this approach could work in patients. Clearly, the answer is yes. Of course, real-world evidence is retrospective and may contain potential biases that we may not recognize, and which could be hard to control. For example, the records come from patients who died in the hospital, accounting for the shorter survivals than often seen. Nonetheless, these data suggest a clinical strategy that should be tested in well-controlled, prospective randomized clinical trials to accelerate the development of a precision medicine strategy for glioblastoma patients.

## Acknowledgements

This work was supported by grants from the National Institute for Neurological Diseases and Stroke (NS73831, NS080939), the Defeat GBM Program of the National Brain Tumor Society, the Ben and Catherine Ivy Foundation, an award from the Sharpe/National Brain Tumor Society Research Program, and a Compute for the Cure Award from the Nvidia Foundation (P.S.M.), as well as the UCSD Neuroscience Microscopy Shared Facility Grant (P30 NS047101). P.S.M. dedicates this paper to Bob Sharpe, a friend and remarkable person who, during his battle with GBM, courageously and gracefully taught me so much about living.

## Author Contributions

J.B. and P.S.M conceived the study and designed all experiments. J.B., J.T., and H.Y. performed experiments. A.K. and A.R. analyzed patient datasets from electronic medical records. S.W., W.Z., R.C.G., and B.P. performed the bioinformatic analysis. J.B., J.T., T.K., S.M., and E.J.C contributed to the mice experiments. A.M.A. and O.Q. performed the lipidomic analysis. H.I.K. and F.B.F. provided cell lines or regents and intellectual input. D.A.W., T.F.C., J.N.R., and B.F.C provided intellectual input. J.B., J.T., S.W., W.Z., A.M.A., O.Q., B.F.C., and P.S.M. analyzed and interpreted data. J.B. and P.S.M. wrote the manuscript, and all authors edited and approved the manuscript.

