## Supplemental information for "Targeting Glioblastoma Signaling and Metabolism with A Re-Purposed Brain-Penetrant Drug"

[1]Department of Pathology, Stanford University School of Medicine, Stanford, CA, USA
[2]ChEM-H, Stanford University, Stanford, CA, USA
[3]Department of Medicine, Institute for Genomics and Systems Biology, University of Chicago, Chicago, IL, USA
[4]Department of Medicine, UCSD School of Medicine, La Jolla, CA, USA
[5]Division of Regenerative Medicine, Department of Medicine, University of California San Diego, La Jolla, CA, USA
[6]Ludwig Institute for Cancer Research, University of California San Diego, La Jolla, CA, USA
[7]Department of Neurosurgery, University of Minnesota, Minneapolis, MN, USA
[8]Department of Pharmacology, UCSD School of Medicine, La Jolla, CA, USA
[9]Department of Neurological Surgery, Northwestern University Feinberg School of Medicine, Chicago, IL, USA.
[10]Department of Pathology, UCSD School of Medicine, La Jolla, CA, USA
[11]Moores Cancer Center, UCSD School of Medicine, La Jolla, CA, USA
[12]Department of Neurology, David Geffen School of Medicine, University of California Los Angeles, CA, USA
[13]Department of Molecular and Medical Pharmacology, David Geffen UCLA School of Medicine, Los Angeles, CA, USA
[14]Jonsson Comprehensive Cancer Center, David Geffen UCLA School of Medicine, Los Angeles, CA, USA
[15]Department of Human Genetics, Institute for Genomics and Systems Biology, University of Chicago, Chicago, IL, USA
[16]Department of Chemistry, The Skaggs Institute for Chemical Biology, The Scripps Research Institute, La Jolla, CA, USA
[17]Lead Contact

### METHODS

#### Cell lines

Patient-derived GBM neurosphere lines were obtained as previously described (Buczkowicz et al., 2014; Laks et al., 2016; Nagaraja et al., 2019; Turner et al., 2017) and were cultured in DMEM/F12 medium supplemented with 1x B27, 20 ng/ ml of EGF, 20 ng/ ml of FGF, 1 µg/ ml heparin and 1x Glutamax (Gibco). U87EGFRvIII cells were established by stably expressing EGFRvIII in U87 cells, as previously described (Wang et al., 2006). U87, U87EGFRvIII, RPE1, and IMR90 cells were maintained in DMEM supplemented with 10% FBS and 1% penicillin/streptomycin, while normal human astrocytes (NHA) were cultured according to the manufacturer's standard protocol by using the astrocyte growth medium BulletKit (Lonza). The attached cells were maintained in 10% FBS medium and changed to 1% FBS medium for follow-up experiments as indicated in the methods. All cell lines were maintained at 37°C in a humidified incubator with 5% $CO_2$. More detailed information for all cell lines used in this study is listed in Table S1.

#### Intracranial GBM xenograft models

All mice experiments were approved by the Institutional Animal Care and Use Committee (IACUC) at the University of California, San Diego. Intracranial GBM xenograft models were established as described previously (Bi et al., 2019). In brief, patient-derived neurosphere cells were first engineered to express a near-infrared fluorescent protein IRFP720, and U87EGFRvIII cells were stably expressed with the turboFP635 protein. A total number of 5 x $10^4$ U87EGFRvIII-FP635 cells, or 5 x $10^4$ GBM39-IRFP cells, or 2.5 x$10^5$ HK296-IRFP cells in 5 µl PBS were intracranially injected into brains of five-week-old female athymic nude mice (Charles River Laboratories). 6~8 mice were injected for each group. For drug treatment, fluoxetine solution stocks were prepared by dissolving fluoxetine in water. Equal volumes of fluoxetine stocks or vehicle (water) were administrated to mice once daily (4.2 mg/kg, 10 mg/kg, 15 mg/kg, or 16.4 mg/kg) by oral gavage after tumors were established at day 8~10. DMSO or temozolomide resuspended in DMSO was administrated to mice once daily (5 mg/kg or 20 mg/kg) via intraperitoneal injection starting from day 8 for 5 days. For U87EGFRvIII xenograft models, 10 mg/kg fluoxetine or vehicle was administrated to mice once daily by oral gavage for 10 days starting from day 8 after tumor cells injection. Tumor growth was assessed using an FMT 2500 fluorescence tomography system (PerkinElmer), and survival dates until the onset of neurologic symptoms were recorded for survival curves. All mice were housed in a conventional barrier facility at 22°C on a 12-hour light/dark cycle with free access to water and food, and their health status was checked by following the protocols.

#### Gene expression and shRNA transduction

Lentivirus *SMPD1* expression plasmid was generated by cloning the full-length coding sequence of *SMPD1* into a pLVX-Puro vector (EcoRI and XbaI). Lentivirus shRNA plasmids were purchased from Sigma, and shRNA sequences are listed in the key resources table. For virus production, lentiviral shRNA or gene expression plasmids were transfected with lentivirus packaging plasmids (Clontech) into HEK293T cells, and the supernatant containing virus was collected at 72 hours after transfection. Virus titers were measured before use, and fresh culture medium was changed to the cells after overnight infection. Infection efficiency and selection concentrations of puromycin were determined for every cell line before the follow-up experiments.

### DepMap data analysis

Gene expression, genetic dependency (combined RNAi-screening from Broad, Novartis, and Marcotte projects), and drug sensitivity (PRISM Repurposing Primary Screen 19Q4) datasets (Corsello et al., 2020; Ghandi et al., 2019; McFarland et al., 2018; Nusinow et al., 2020) of Cancer Cell Line Encyclopedia (CCLE) were downloaded from the DepMap portal (https://depmap.org/portal). Mean differences of mRNA expression levels and shRNA dependency scores (DEMETER2) were calculated between glioma cell lines and other CCLE cell lines, and two-tailed Student's t-test was performed to test the significance. For drug sensitivity, the cell viability response to selected compounds (replicate collapsed log fold change values relative to DMSO) of all glioma cell lines from the DepMap dataset was analyzed by Pearson correlation and plotted as a drug-drug sensitivity matrix.

### Sphingolipid analysis by LC-MS

GBM cells were treated with DMSO or 5 µM fluoxetine in 1% FBS DMEM medium for 42 hours and harvested as pellets. Samples were then spiked with a set of internal standards and extracted with an organic solvent system consisting of equal parts of dichloromethane and methanol. Phase separation was achieved by the addition of an equal part of water. The organic layer was collected, the solvent was removed under argon, and the samples were reconstituted in 100 ul of isopropanol/acetonitrile (40/60, v/v). Sphingolipids were separated by liquid chromatography (LC), and the eluting metabolites were measured by mass spectrometry (MS/MS) according to methodologies established at the UCSD LIPID MAPS Lipidomics Core (http://www.ucsd-lipidmaps.org). Briefly, a Waters Acquity UPLC system (Waters Technologies, Milford, MA) with a Phenomenex Kinetex C18 column,150x2.1 mm, 1.7 µm (Phenomenex, Torrance, CA) was used for chromatographic separation. Gradient elution started at 40 % mobile phase B for 10 minutes, then increased linearly to 100 % B over 10 minutes, kept at 100 % B for 30 minutes, and the column was equilibrated with 40 % B for 8 minutes. Buffer A consisted of 100% H2O with 10mM ammonium formate and 0.1% formic acid modifiers. Buffer B consisted of isopropanol/acetonitrile (40/60, v/v) with 10mM ammonium formate and 0.1% formic acid as modifiers. The flow rate was 300 µl/minute, and 10 µl of sample was injected via autosampler.

The LC eluent was interfaced with a mass spectrometer 6500 QTrap (Sciex, Framingham, MA), controlled by Analyst v. 1.7 software, operated in Information Dependent Acquisition mode (IDA), using an Enhanced MS (EMS) scan from m/z 400-1000 at 10000 Da/s as a survey scan. Source parameters were automatically optimized using flow injection analysis into an isocratic flow of 80 % mobile phase B using individual lipid molecules. To maximize metabolite coverage and identification, the sphingolipids were analyzed in positive and negative ion modes. The optimized source parameters of the Turbo V ion source for positive ion mode were as follows: Curtain Gas, 20; Collision Gas, High; IonSpray voltage, 5000; Temperature, 300; Gas 1, 30; Gas 2, 30, Declustering Potential, 100; Collision Energy Spread, 0; Collision Energy, 45. Source parameters for negative ion mode were: Curtain Gas, 20; Collision Gas, High; IonSpray voltage, -4500; Temperature, 300; Gas 1, 30; Gas 2, 30, Declustering Potential, -150; Collision Energy Spread, 0; Collision Energy, -10. From each survey scan, the ions exceeding a pre-set intensity threshold were chosen for Enhanced Product Ion scans (EPI). The lipid molecules were identified by molecular mass, elution time and MS/MS fragmentation patterns. At least three biological replicates were performed for each cell line per treatment. For quantitation, the MS signals of endogenous sphingolipid molecules were normalized to that of the internal standards and the cell numbers of each sample.

#### SMPD1 enzymatic activity assay

The enzymatic activity of acid sphingomyelinase (SMPD1/ASM) was measured by the cleavage of HMU-PC using a commercial kit (Echelon, K-3200) as described previously (van Diggelen et al., 2005). GBM cells with indicated hours DMSO or fluoxetine treatment were collected. Cell pellets were then resuspended in water with proteinase inhibitor and sonicated in an ice water bath for 10 cycles (30 seconds on and 30 seconds off). Tumor samples were first homogenized in water with proteinase inhibitor by a rotor-stator tissue homogenizer and then sonicated on ice. After 5-minutes centrifugation (10,000 x g) at 4˚C, protein concentration of each sample was determined. Equal volumes of 10 μg samples were added into each reaction and incubated at 37°C for 3 hours. After adding the stop buffer, the fluorescence of HMU was recorded on an Infinite M1000 Plate Reader (Tecan) at 360 nm excitation and 460 nm emission. Data were normalized to that of the indicated control group and plotted from four biological replicates.

#### Cell viability assay

Cell viability was assessed using a CellTiter-Glo luminescent cell viability assay kit (Promega). Attached cells or GBM neurosphere cells were seeded into each well of 384-well plates with DMEM medium supplemented with 1% FBS and 1% penicillin/streptomycin or DMEM/F12 medium supplemented with 1/4x B27, 20 ng/ ml of EGF, 20 ng/ ml of FGF, 1 μg/ ml heparin, and 1x Glutamax respectively. Equal volumes of vehicles or drugs diluted with the medium were added into the wells the next day, and the cells were cultured for 72 hours. After 15 minutes of incubation with CellTiter-Glo reagent at room temperature, the luminescent was recorded using an Infinite M1000 Plate Reader (Tecan). Four biological replicates were performed for each cell line per treatment. The area under the curve (AUC) was calculated by the AUC function in the DescTools R package with the “spline” method, which results in the area under the natural cubic spline interpolation.

#### Cell death and Annexin V-positive cell analysis

Annexin V-positive cells were determined by flow cytometry using a FITC Annexin V Apoptosis Detection Kit (BD Biosciences). In brief, cells were treated with DMSO or fluoxetine and cultured for 72 hours in DMEM medium with 1% FBS (attached cells) or DMEM/F12 medium supplemented with 1/4x B27, 20 ng/ ml of EGF, 20 ng/ ml of FGF, 1 µg/ ml heparin and 1x Glutamax (neurosphere cells). For shRNA experiments, cells after shRNA lentivirus infection were reseeded and cultured for 72 hours. Cells were then collected for Annexin V/ PI staining and analyzed by using a BD LSRII flow cytometer (BD Biosciences). For cell death trypan blue assay, cells were seeded in 6-well plates (attached cells) or 25 $cm^2$ flasks (neurosphere cells) and cultured for five days after shRNA lentivirus infection or with drug treatment. Dead cells and live cells were counted by trypan blue assay using a TC10 automatic cell counter (Bio-Rad). At least three biological replicates were performed for each cell line per treatment.

#### Soft-agar colony formation assay

For each well of 12-well plates, 2000 GBM cells were mixed with 0.4% low-gelling-temperature agarose (Sigma) in growth medium and immediately plated onto a solidified bottom layer containing 1% low-gelling-temperature agarose in the growth medium. Four biological replicates were performed for each treatment, and Cells were treated and fed with fresh growth medium every three days for 3 weeks. Colonies were then stained with 0.005% crystal violet, imaged by a ChemiDoc MP imaging system (Bio-Rad), and counted by ImageJ.

#### Crystal violet clonogenic assay

24 hours after shRNA or gene expression lentivirus infection, 2000 GBM cells were reseeded into each well of 6-well plates and cultured in 2 ml growth medium for 2 weeks. The medium was refreshed every three days and removed before crystal violet staining. Colonies were fixed with 80% methanol in $ddH_2O$ for 10 minutes and stained with 0.05% crystal violet for 20 minutes. After washed with water, plates were imaged on a ChemiDoc MP imaging system (Bio-Rad), and colony density was quantified by ImageJ.

#### RNA extraction and qRT-PCR

Total RNA extraction was performed using an RNeasy mini kit (QIAGEN) and the SuperScript IV VILO master mix (Invitrogen) was used for reverse transcription. Samples are mixed with primers and SYBR Green Supermix and amplified on a CFX96 real-time PCR detection system (Bio-Rad). The results were processed by the $\Delta\Delta C_t$ method, and expression levels were normalized to the reference gene and indicated control group.

#### Immunofluorescence staining

Cells were seeded in laminin-coated chamber slides and treated with DMSO or 5 µM fluoxetine for 42 hours. After washed twice with PBS, cells were fixed in 4% PFA for 15 minutes, permeabilized with 0.2% Triton X-100 in PBS for 15 minutes and then blocked with 2% BSA in PBS for 45 minutes. Primary antibodies (anti-LAMP1, #9091, Cell Signaling at 1: 200 dilution; anti-phospho-histone H2A.X (Ser139), 05-636, Millipore at 1:200 dilution) in PBS with 0.02% Triton X-100 and 0.5% BSA was applied to cells and incubated overnight at 4°C. After four washes with PBS, cells were then incubated with fluorescent secondary antibody Alexa Fluor anti-Rabbit 546 (A11010, Invitrogen) or Alexa Fluor anti-Mouse 488 (A11017, Invitrogen) at a dilution of 1:1000 in PBS at room temperature for 1 hour. For EGFR staining, an EGFR antibody conjugated with Alexa Fluor 488 (#5616, Cell Signaling) was used at 1:200 dilution in overnight incubation. For LysoTracker staining, after 42 hours treatment of DMSO or 5 μM fluoxetine, cells were washed with PBS and incubated with 50 nM LysoTracker (L7528, Invitrogen) at 37 °C for 1 hour. Cells were then washed with PBS, fixed in 4% PFA for 15 minutes. After four washes with PBS, cells were mounted with antifade reagent with DAPI (Life Technologies) for imaging on an Olympus FV1000 confocal microscope. Fluorescent intensity was quantified by ImageJ.

### Drug and lipid treatment

For western blot, cells were collected after 48 hours of treatment unless otherwise indicated. GBM neurosphere cells were seeded in DMEM/F12 medium supplemented with 1/4x B27, 20 ng/ ml of EGF, 20 ng/ ml of FGF, 1 μg/ ml heparin and 1x Glutamax, cultured overnight, and treated with DMSO or fluoxetine for indicated hours before harvesting for enzymatic assay or western blots. In proteasome and lysosome inhibitors experiments, cells were treated with DMSO or fluoxetine for 42 hours and incubated for additional 6 hours in the absence or presence of 10 μM MG132 or 50 μM Chloroquine before collecting. For lipid treatment, 10 μM of sphingolipids conjugated with BSA or an equal amount of BSA solution was added into the growth medium and mixed on a shaker at room temperature for 30 minutes before treating cells. Cells were then cultured with the lipid or vehicle-adding medium for 48 hours and applied for further western blot analysis or immunofluorescence staining. In cell viability assay, GBM cells were firstly cultured with lipid or vehicle -adding medium overnight and then incubated with fluoxetine or DMSO for 72 hours before CellTiter-Glo assay.

### Membrane lipid order imaging of live cells

Membrane lipid order imaging of live GBM cells was performed as described previously (Owen et al., 2011). Briefly, GBM cells in glass-bottom dishes were treated with 5 μM fluoxetine for 40 hours in 1% FBS DMEM medium and then stained with 5 μM Laurdan (D250, Invitrogen) for 3 hours in serum-free medium at 37°C in a humidified incubator with 5% CO2. Cells were then imaged on a Leica SP5 Confocal/Multiphoton system with the excitation at 800 nm and the emission at 400–460 nm and 470–530 nm). Pseudocolored generalized polarization (GP) images were achieved by using an ImageJ plug-in as described (Owen et al., 2011). GP values at the plasma membrane region of at least 60 cells were quantified by ImageJ and plotted as histograms.

### Western blot analysis

Cells were washed with cold PBS and lysed with 1x RIPA lysis buffer containing 1x protease and phosphatase inhibitor cocktail on ice for 30 minutes. Tumor samples were homogenized on ice with cold PBS supplemented with protease and phosphatase inhibitor cocktail and then lysed with an equal volume of 2x RIPA buffer on ice for 30 minutes. BCA protein assay kit (Thermo Scientific) was used to determine the protein concentration. Equal amounts of protein samples were mixed with Laemmli sample buffer, boiled at 100°C for 5 minutes, electrophoresed using 4%–12% NuPAGE Bis-Tris mini gels, and then transferred onto nitrocellulose membranes by a Trans-Blot Turbo transfer system (Bio-Rad). Membranes were blocked with 5% BSA in TBST buffer and incubated with corresponding primary antibodies at 4°C overnight, followed by incubation with HRP-conjugated secondary antibodies at room temperature for 1 hour. After washing, the blots were developed with SuperSignal West Pico chemiluminescent substrate (Thermo Scientific) and imaged using Image Lab software on a ChemiDoc MP imaging system (Bio-Rad).

### Immunohistochemistry analysis

Formalin-fixed, paraffin-embedded tissue sections were performed by the Tissue Technology Shared Resource (TTSR)-Histology Core at UCSD. Standard staining protocols were followed. In brief, the antigen was retrieved by boiling slides in 0.01 M of sodium citrate (pH 6.0) for 15 minutes. Tissue sections were then incubated with primary antibodies overnight at 4°C, followed with 30 minutes incubation with biotinylated secondary antibodies at room temperature. Tissue sections for TUNEL staining were incubated with TdT/ dUTP at 37°C for 30 minutes and then with HRP-conjugated anti-digoxigenin at room temperature for another 30 minutes. Stained slides were imaged on an Olympus BX43 microscope and quantified in a double-blind fashion using Visiopharm image analysis software.

### Cell surface EGFR and internalization analysis

The intensity of cell surface EGFR was determined by flow cytometry with an EGFR antibody that recognizes the extracellular domains of both wild-type EGFR and EGFRvIII proteins as described previously (Lu et al., 2007; Luwor et al., 2001). In brief, GBM39 and HK296 cells were first chilled on ice for 20 minutes and then incubated with a primary EGFR antibody (GR01, Millipore, mAb528, 1:20) on ice for one hour. After gently washed once with 10 ml cold PBS, the cells were incubated with an Alexa Fluor 488 goat anti-mouse second antibody (A11017, Invitrogen, 1:500) on ice for an additional hour. The cells were then gently washed once with 10 ml cold PBS and analyzed on a BD LSRFortessa X-20 flow cytometer (BD Biosciences). For EGFR internalization assay, GBM39 cells treated with 5 μM fluoxetine or DMSO for 48 hours were chilled on ice and incubated with the EGFR antibody (GR01, Millipore, mAb528, 1:40) on ice for one hour. Primary antibody-stained cells were either kept on ice or moved to 37 ℃ for 15 or 30 minutes to allow internalization. Following internalization, the cells were washed and incubated with Alexa Fluor 488 goat anti-mouse second antibody (Cat#A11017, Invitrogen, 1:500) for another hour before analyzing by flow cytometer. Internalized EGFR level was defined as the decreased signal of surface EGFR after incubation at 37 ℃.

Three to four biological replicates were performed for each treatment. A total of 10, 000 events for each sample was recorded and analyzed.

### Density gradient fractionation

The detergent-free density gradient fractionation was performed as previously described (Cizmecioglu et al., 2016; Macdonald and Pike, 2005). In brief, GBM39 cells treated with 5 µM fluoxetine or DMSO for 48 hours were pelleted at 250 g for 5 minutes, washed once with cold PBS, and resuspended in 1 ml of cold homogenization buffer (20 mM Tris-HCl, pH7.8, 0.250 M sucrose, 1 mM $CaCl_2$ and 1 mM $MgCl_2$) with protease and phosphatase inhibitors. Homogenates were then passed through a 23 g needle for 20 times followed by centrifugation at 4°C at 1000 g for 10 min. 1 ml of supernatants were collected, mixed with 1 ml of 50% Opti-Prep solution (Sigma), and placed in the bottom of a 5 ml Ultra-Clear centrifuge tube (Beckman Coulter). 400 µl each of 20%, 17.5%, 15%, 12.5%, 10%, 7.5% and 5% Opti-Prep solutions were then poured onto the top. After ultracentrifugation at 100, 000 g for 2 hours at 4°C using an SW-55Ti rotor in a Beckman ultracentrifuge, equal volumes of six fractions were collected from the top layer to bottom layer and loaded for further western blot analysis. Lipid rafts fractions were characterized by the non-lipid rafts marker Calnexin and lipid rafts markers Flotilin-1 and Gα(q). The percentages of protein level in fraction 1 were calculated by dividing the amount of protein in fraction 1 by the total amount of protein in all six fractions and plotted from three independent experiments.

### RNA-seq analysis

Neurosphere cells were treated with DMSO or 5 µM fluoxetine for 42 hours and collected for RNA extraction. RNA sequencing was performed by Novogene. RNA-Seq reads were aligned to the human reference transcriptome (GRCh38 release-98) and quantified using the Salmon software (Patro et al., 2017). The --gcBias flag was used to estimate a correction factor for systematic biases commonly present in RNA-seq data. The differential expression analysis was performed using the likelihood ratio test (LRT) in DESeq2(Love et al., 2014). The LRT examines two nested models for the read counts, a full model where gene expression was explained by fluoxetine treatment and cell lines and a reduced model, in which only cell lines were considered. The test determines if fluoxetine treatment contributed significantly to the gene expression beyond the expected expression level due to cell lines. Gene Set Enrichment Analysis (GSEA) was performed on all genes ranked by likelihood ratio test statistic against MSigDB v7.1 (Subramanian et al., 2005). Enriched terms, including EGFR signaling inhibitor down and up signatures (Kobayashi et al., 2006) (https://www.gsea-msigdb.org/gsea/msigdb/cards/KOBAYASHI_EGFR_SIGNALING_24HR_DN and https://www.gsea-msigdb.org/gsea/msigdb/geneset_page.jsp?geneSetName=KOBAYASHI_EGFR_SIGNALING_24HR_UP), were visualized using ClueGO (Bindea et al., 2009).

### TCGA data analysis

TCGA GBM RNA-seq dataset was used in patient survival study. In survival association group analysis, we used the “lifelines” package in python to fit Cox proportional hazard models (Andersen, 1982). *P* values were calculated by log likelihood ratio tests. To evaluate whether a gene’s expression provides additional prognostic information beyond the baseline survival probability due to age at diagnosis, we compared the likelihood of two nested models: a full model with gene expression and age of patients and a reduced model, in which only age was considered. Proportional hazards model and log-rank test were applied to assess the prognostic significance of individual genes. Overall survival of GBM patients with the top 25% and bottom 25% of *SMPD1* expression in the TCGA GBM cohort (RNA-seq) was statistically compared by Log-rank test. Cox proportional hazard ratios were calculated. *P* values and numbers of patients of each cohort were indicated in the figure. The Gene Expression Profiling Interactive Analysis (GEPIA) web-server(Tang et al., 2017) was used to analyze the expression of metabolic genes between GBM tumors and normal brains based on TCGA and GTEx RNA-seq data. The genetic alterations of *EGFR*, *SMPD1*, and *IDH1* in the merged cohort of LGG and GBM TCGA (PanCancer Altas) datasets were assessed using cBioPortal for Cancer Genomics (http://www.cbioportal.org) (Cerami et al., 2012; Gao et al., 2013). Gene set enrichment analysis was performed to characterize genes differentially expressed in GBM clinical samples (TCGA GBM, HGU133A) with high or low *SMPD1* expression. The median of the GBM cohort was chosen as the cutoff for high and low *SMPD1* expression groups.

#### Evaluation of Fluoxetine in Overall Survival of GBM Patients in Electronic Medical Records

Complete details of the database used, data overview, use of ICD9 and ICD10 codes, data extraction pipeline, exclusion criteria, enrichment for GBM patients, statistical analyses, and full results are presented in Supplemental Document S1.

### QUANTIFICATION AND STATISTICAL ANALYSIS

Unless otherwise indicated, all statistical analysis was performed using GraphPad Prism 8 software. Unpaired two-tailed Student’s t-test was applied to compare two experimental groups, and one-way or two-way ANOVA followed by multiple comparisons test was used to assess differences between three or more experimental groups. Paired two-tailed Student’s t-test was only performed in comparing the percentage of protein intensity in paired samples. Log-rank (Mantel-Cox) test was used in survival analysis. Fisher’s exact test was applied for mutual exclusivity analysis of genetic alterations in TCGA clinical samples and tumor recurrence analysis of PDX mice models. Bar graphs show mean ± SD or SEM as indicated in the legends, and p values less than 0.05 are defined to be significant. Numbers of samples, statistical tests, and p values analyzed in each experiment are reported in the respective figure legends or methods.

### SUPPLEMENTAL ITEM TITLES AND LEGENDS

**Document S1.** Evaluation of Fluoxetine in Overall Survival of GBM Patients in Electronic Medical Records from The IBM Marketscan Dataset.

**Table S1.** Information of Cell Lines Used. It includes major clinical and genomic features, such as age, gender, tumor category, and prior treatments, of 18 patient-derived adult GBM and DIPG neurosphere lines that were utilized in this study.

### Supplemental Figures

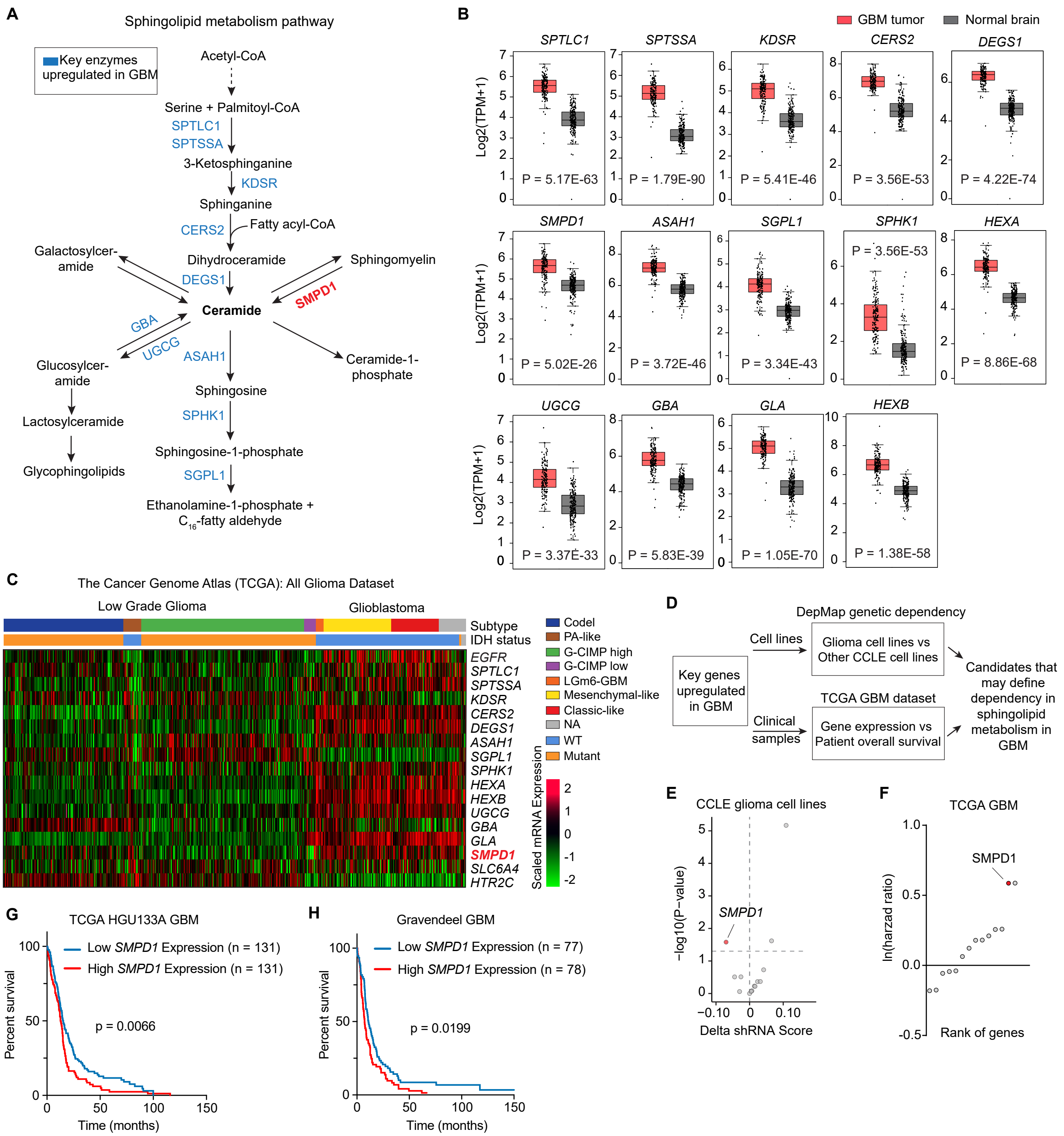


**Figure S1. GBMs Are Highly Dependent on Sphingolipid Metabolism.** (A) Schematic pathways of sphingolipid metabolism. (B) 14 coding genes of key enzymes in the sphingolipid metabolism pathway are highly expressed in GBM tumors (TCGA) in comparing with normal brains (GTEX). (C) Heatmap of gene expression in the merged cohort of low-grade glioma and glioblastoma of TCGA datasets. (D) Outline of the strategy utilized for identifying metabolic dependency of GBM in the sphingolipid pathway. (E) Gene dependency analysis of 14 upregulated genes (B) in the sphingolipid metabolism pathway from the DepMap dataset. Glioma cells show a much more significant dependency on *SMPD1* than other sphingolipid metabolic genes. (F) Rank of genes with the association (hazard ratio) between mRNA expression and overall survival of GBM patients from the TCGA dataset. High *SMPD1* expression is significantly correlated with poor overall survival of GBM patients. (G and H) Kaplan-Meier analysis of overall survival in GBM patients with high or low *SMPD1* mRNA expression in the TCGA HGU133A GBM and Gravendeel GBM cohorts. Two-tailed Student's t-test for (B) and (E). Log-rank test for (G) and (H). The median value (center line), the min and max (whiskers), and the 25th and 75th percentiles (box perimeters) are presented in (B).

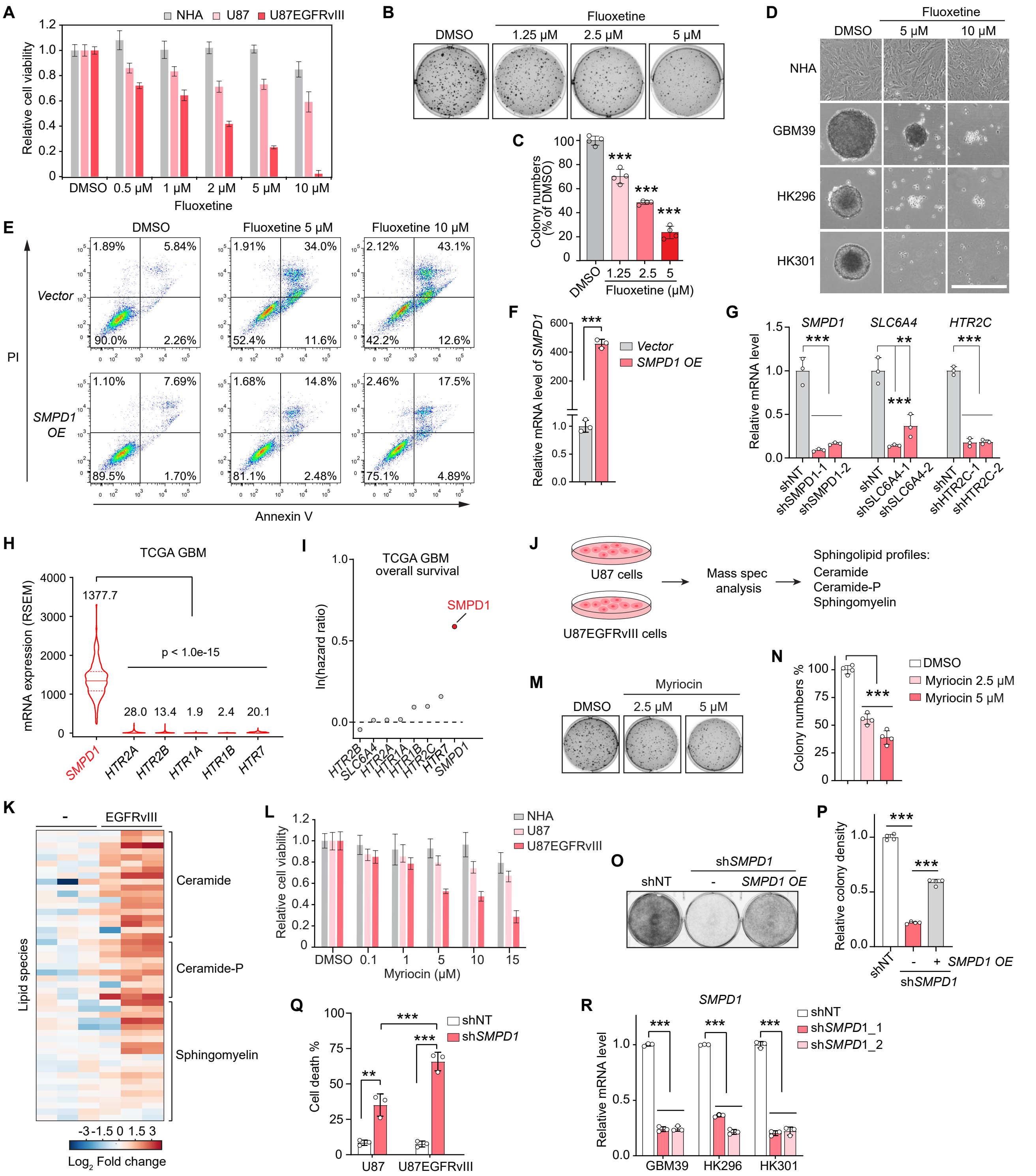

**Figure S2. Fluoxetine Selectively Kills GBM Cells through SMPD1 Inhibition.** (A) Relative cell viability of normal human astrocytes (NHA), U87, and U87 cells expressing *EGFRvIII* treated by fluoxetine (n= 4). (B and C) Colony formation of U87EGFRvIII cells in soft agar (n= 4). (D) Representative images of NHA and GBM spheres after DMSO or fluoxetine treatment. Scale bar, 500 μm. (E) Representative flow cytometry analysis of Propidium iodide (PI) and Annexin V positive cells after DMSO or fluoxetine treatment. (F) Relative mRNA level of *SMPD1* with its overexpression in (E). (G) shRNA knockdown efficiency of *SMPD1*, *SCL6A4*, and *HTR2C* in U87 cells expressing EGFRvIII. (H) mRNA level (RSEM) in GBM patient samples (TCGA). (I) Rank of genes with the association (hazard ratio) between mRNA expression and overall survival of GBM patients from the TCGA dataset. (J and K) Lipidomic analysis of U87 cells with or without EGFRvIII expression (n= 3). (L-N) Relative cell viability (L) and colony formation in soft agar (M and N) in NHA and GBM cells treated with myriocin, an inhibitor of sphingolipid *de novo* synthesis. (O and P) Crystal violet staining for U87 cells expressing EGFRvIII (n= 4). (Q) Percentage of cell death of U87 cells with or without *EGFRvIII* overexpression after *SMPD1* or non-targeting shRNA transfection (n= 3). (R) Relative mRNA level of *SMPD1* in three patient-derived GBM lines with indicated shRNA transfection. For (F), a two-tailed Student's t-test was performed. ANOVA followed by Tukey's multiple comparisons test was applied in (C), (G), (H), (N), and (P-R). Data represent mean± SD except (H). The median value (center line) and the 25th and 75th percentiles (dash lines) are presented in (H). **p < 0.01; ***p < 0.001.

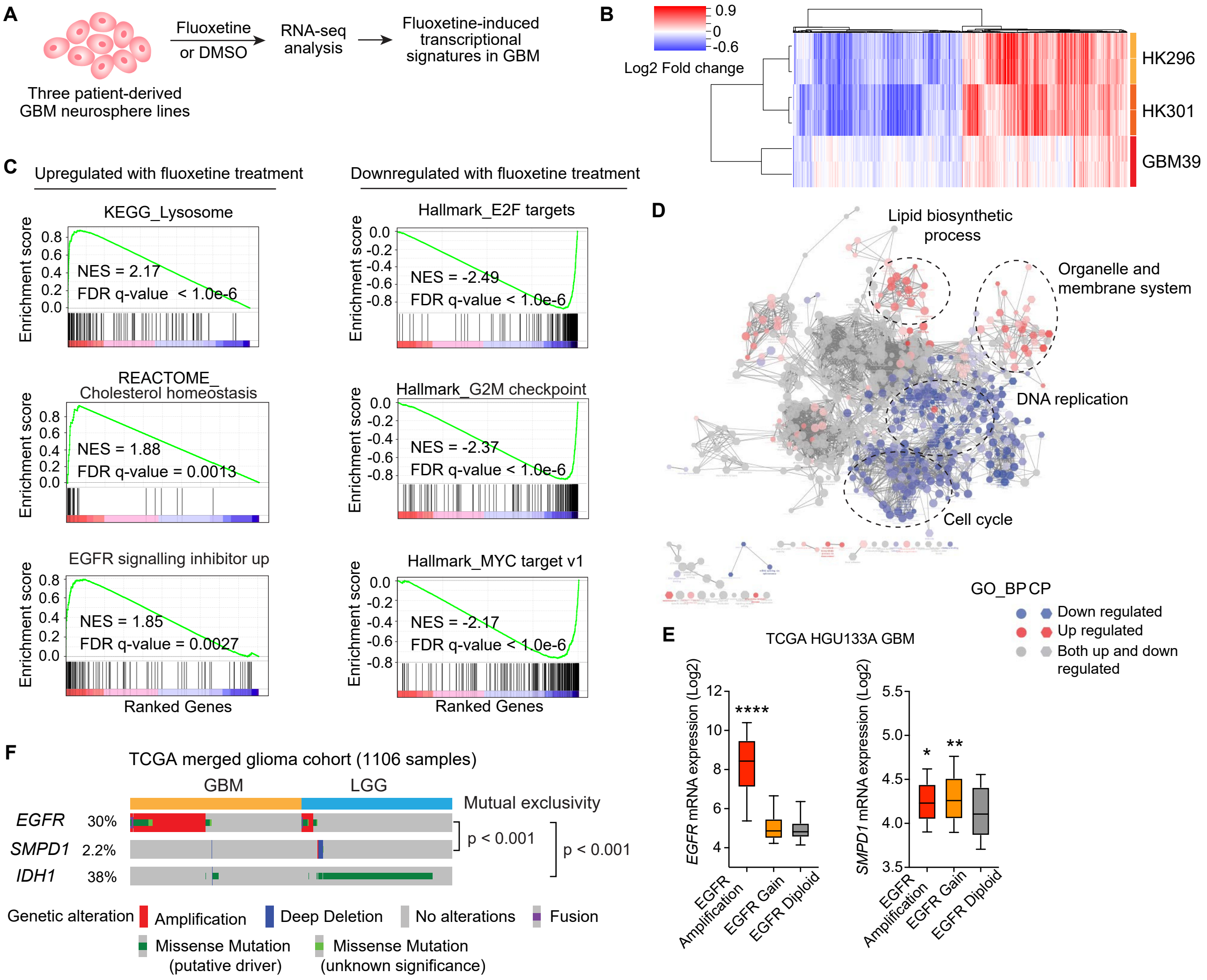


**Figure S3. Fluoxetine-Induced Transcriptional Signatures in GBM Cells.** (A) Schematic overview of RNA-seq analysis in 3 GBM neurosphere lines with 42 hours DMSO or fluoxetine treatment. Two biological replicates per treatment were performed for each cell line. (B) Heatmap of differentially expressed genes (FDR < 0.05) between fluoxetine and DMSO treatment in GBM39, HK296, and HK301 cells. (C) Selected most significantly upregulated or downregulated transcriptional signatures by GSEA. FDR q-value and NES are shown. (D) Gene Ontology (GO) enrichment analysis showing both upregulated and downregulated transcriptional signature clusters. (E) *EGFR* and *SMPD1* mRNA level in GBM clinical samples of TCGA GBM (HUG133A) cohort. (F) Genetic alterations of *EGFR*, *SMPD1*, and *IDH1* in merged LGG and GBM cohort of TCGA dataset (1106 samples). Deep deletion of *SMPD1* is mutually exclusively associated with *EGFR* amplification and gain of function mutations in clinical tumor samples. For (E), the median value (center line), the min and max (whiskers), and the 25th and 75th percentiles (box perimeters) are presented. Fisher's exact test for (F), and ANOVA followed by Tukey's multiple comparisons test for (E). $*p < 0.05$; $**p < 0.01$; $****p < 0.0001$.

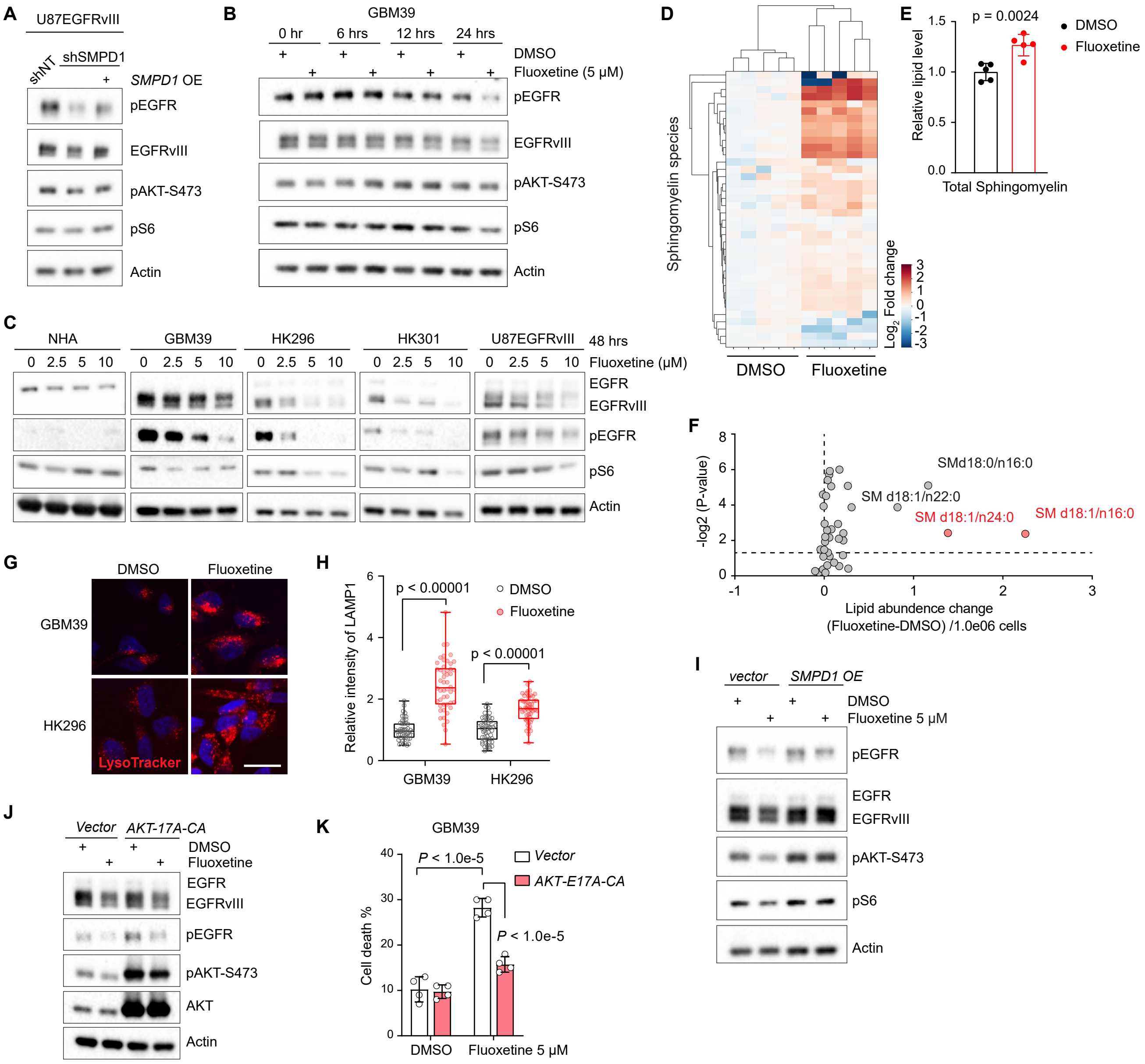


**Figure S4. Fluoxetine Acts as an Indirect Inhibitor of EGFRvIII by Blocking SMPD1 and Elevating Sphingomyelin Levels.** (A) Western blot analysis of EGFR signaling activity in GBM cells expressing non-targeting (NT) or *SMPD1* shRNA. (B) Western blot analysis of EGFR signaling activity in GBM39 cells treated with DMSO or 5 μM fluoxetine for indicated hours. (C) Western blot analysis of EGFR signaling activity in normal human astrocytes (NHA) and GBM cells after 48 hours of fluoxetine treatment. (D-F) Sphingomyelin profiles identified by mass spectrometry in U87EGFRvIII cells with DMSO or fluoxetine treatment (n=5). SM d18:1/n16:0 and SM d18:1/n24:0 are two sphingomyelin species that were dramatically elevated after fluoxetine treatment. (G) Lysotracker staining in GBM39 and HK296 cells. Scale bar, 20 μm. (H) Quantification of LAMP1 intensity in GBM39 and HK296 cells (n= 50). (I) Western blot showing EGFR and downstream signaling in U87EGFRvIII cells with indicated treatment and gene expression. (J and K) AKT signaling and percentage of cell death in GBM39 cells expressed vector or a constitutively active AKT E17A-CA allele (n= 4). Data represent mean± SD in (E). The median value (center line), the min and max (whiskers), and the 25th and 75th percentiles (box perimeters) are presented in (H). Two-tailed Student's t-test for (E), (F), and (H). ANOVA followed by Tukey's multiple comparisons test for (K).

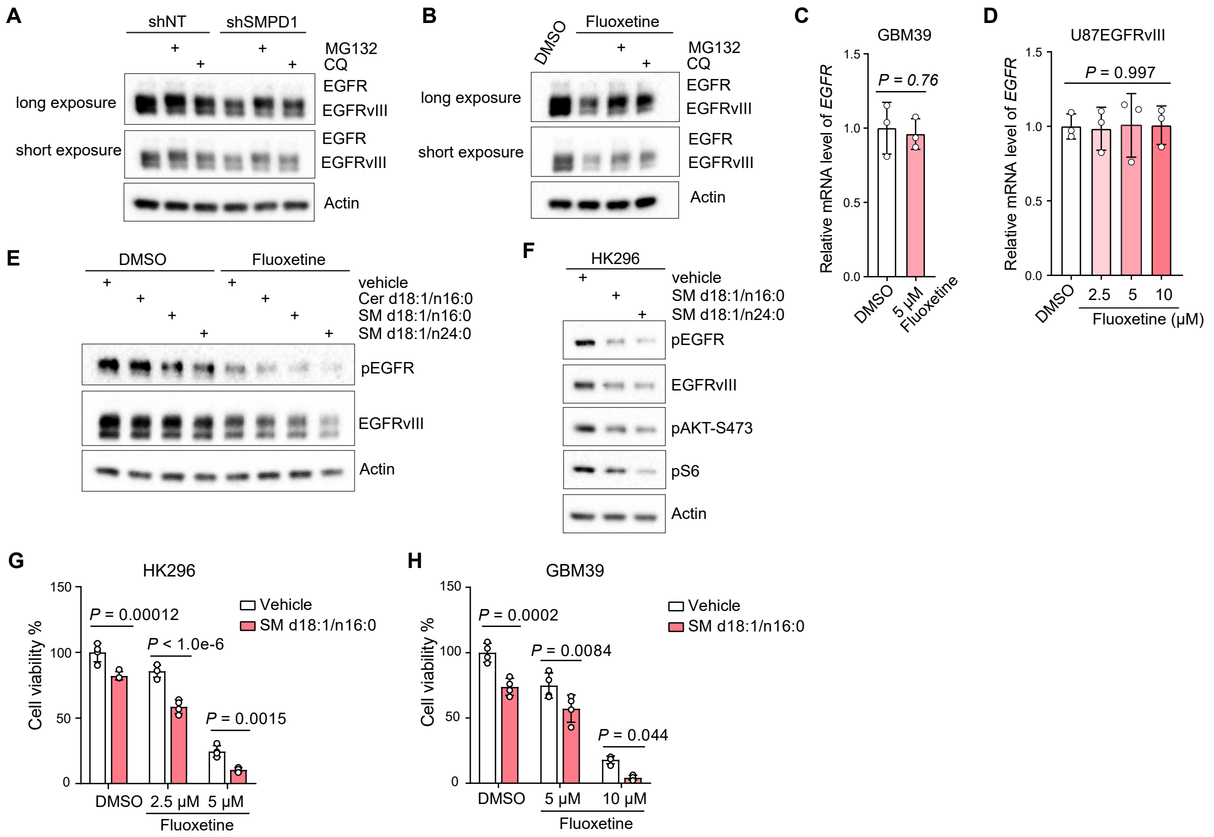


**Figure S5. Fluoxetine Kills GBM Cells by Inhibiting SMPD1 and Abrogating EGFRvIII Signaling.** (A and B) Western blot analysis of EGFR protein in GBM cells with indicated treatment and further incubated for additional 6 hours in the absence or presence of MG132 or Chloroquine (CQ). (C and D) Relative *EGFR* mRNA level in GBM cells with 48 hours of fluoxetine treatments (n= 3). (E-H) Adding sphingomyelins, but not ceramides, suppresses EGFR signaling (E and F) and further decreases cell viability (G and H) in GBM cells (n= 4). Data represent mean± SD. Two-tailed Student's t-test for (C). ANOVA followed by Tukey's multiple comparisons test for (D), (G), and (H).

A

| FDA-approved fluoxetine doses for human non-cancer indications (mg/day) | Mice doses converted from human doses based on body surface area (mg/kg/day) |
|---|---|
| 20 | 4.2 |
| 40 | 8.2 |
| 50 | 10.2 |
| 60 | 12.3 |
| 80 | 16.4 |

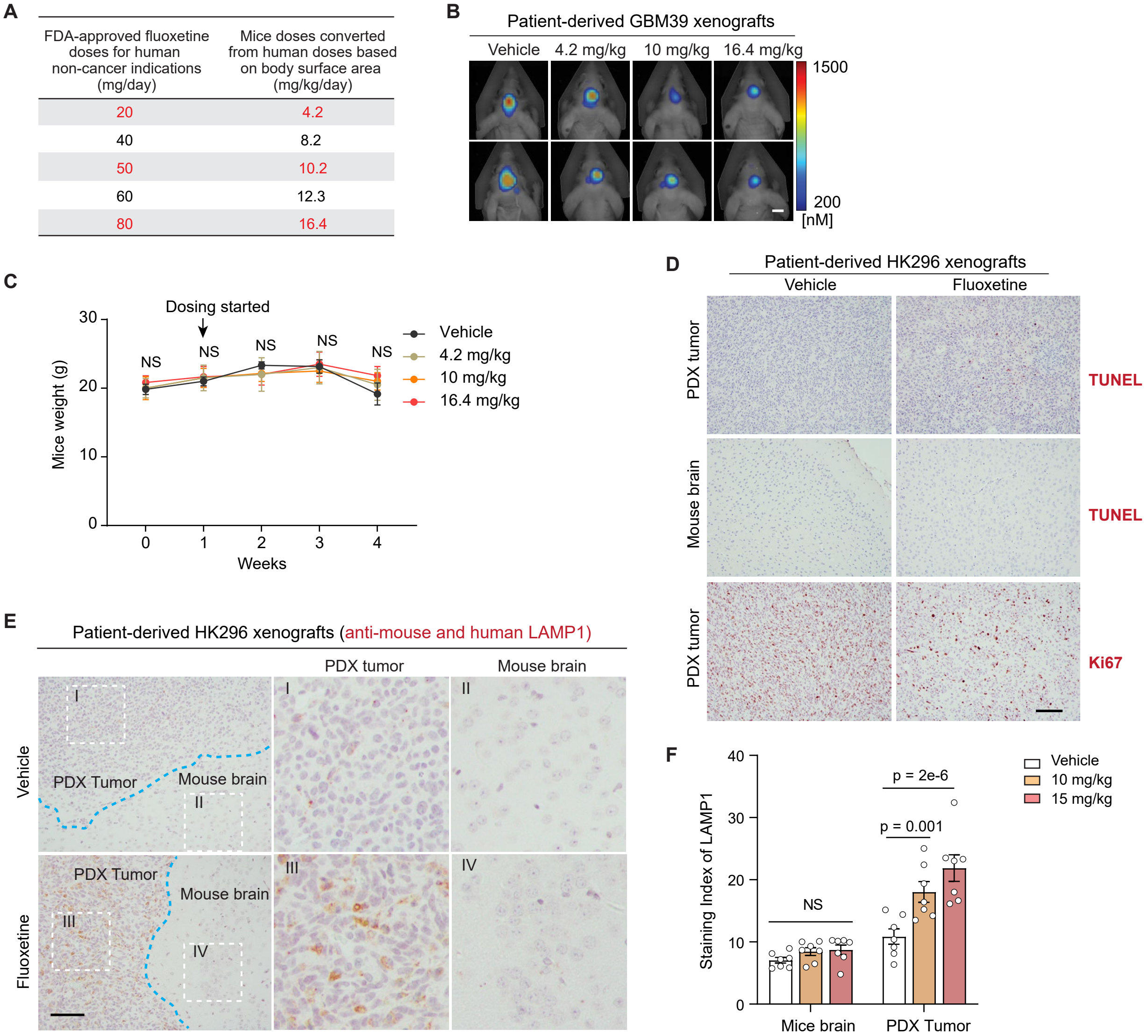


**Figure S6. Fluoxetine Suppresses Tumor Growth in Patient-Derived Orthotopic GBM Models.** (A) FDA-approved doses of fluoxetine for human non-cancer indications and converted mice doses based on body surface area. (B) Representative tumor images of GBM39 patient-derived orthotopic mice models. Scale bar, 5 mm. (C) Body weights of mice bearing patient-derived GBM39 xenograft models with indicated administrations (n= 6 per group). Data represent mean± SD. (D) Representative images of immunohistochemistry analysis of TUNEL and Ki67 in HK296 xenograft tumors and surrounding mice brains. Scale bar, 100 µm. (E and F) Immunohistochemistry analysis of LAMP1 in HK296 xenograft tumors and surrounding mice brains by using an antibody that could detect both human and mice LAMP1 proteins (n= 7). Scale bar, 100 µm. Data represent mean± SEM. ANOVA followed by Tukey's multiple comparisons test in (C) and (F). NS, not significant.

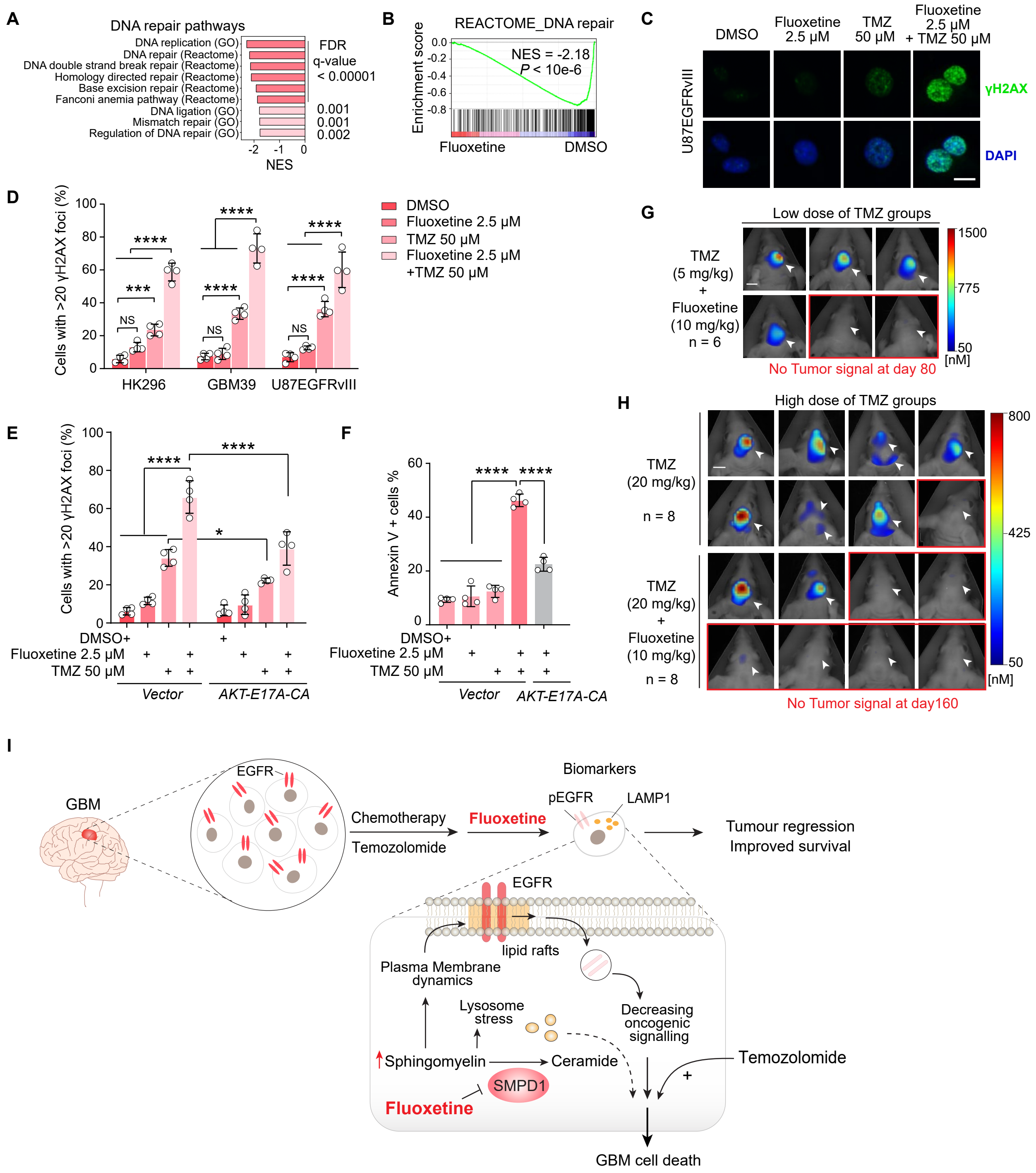

**Figure S7. Combination of Fluoxetine and Temozolomide Induces DNA Damage and Suppresses GBM Tumor Recurrence.** (A and B) Downregulated DNA repair pathways in three patient-derived GBM neurosphere lines after fluoxetine treatment. (C-E) γH2AX staining analysis of GBM cells. Data are from four replicate samples for each treatment. Scale bar, 10 μm. (F) Percentage of Annexin V-positive GBM cells with indicated treatment (n= 4). (G and H) Tumor images of mice with the combination treatment of fluoxetine and TMZ at day 80 or day 160 in GBM39 xenograft models. No tumor signal was identified in 2 mice of low dose TMZ (5 mg/kg) combination group and 6 mice of high dose TMZ (20 mg/kg) combination group. Arrowheads indicate the injection sites of tumor cells. Scale bar, 5 mm. (I) Schematic model of the fluoxetine's anti-GBM effect through SMPD1 inhibition. GBMs are highly dependent on SMPD1 for survival, which is enhanced in *EGFRvIII*-amplified tumors. Fluoxetine kills GBM cells by indirectly inhibiting EGFRvIII by elevating sphingomyelin levels to alter plasma membrane organization, and by causing lysosomal stress. Fluoxetine can be combined with temozolomide to suppress tumor recurrence, as detected in preclinical *in vivo* models and in electronic medical record data from patients. Data represent mean± SD. ANOVA followed by Tukey's multiple comparisons test in (D-F). *p < 0.05; ***p < 0.001; ****p < 0.0001; NS, not significant.

### Table S1. Information of Cell Lines Used.

| Cell line name | Cell type | Molecular status | Other major genetic alterations | Disease | Age | Gender | Tumor category | Prior treatment | Source reference |
|---|---|---|---|---|---|---|---|---|---|
| RPE1 | Human epithelial cells immortalized with hTERT | - | hTERT | Normal | n/a | Female | - | n/a | ATCC |
| NHA | Normal Human Astrocytes | - | - | Normal | n/a | n/a | - | n/a | Lonza |
| IMR90 | Human lung fibroblast | - | - | Normal | 16 weeks | Female | - | n/a | ATCC |
| U87EGFRvIII | Human GBM cells | EGFRvIII overexpression | PTEN loss, CDKN2A loss | GBM | n/a | Male | n/a | n/a | Wang et al., 2006 |
| GBM39 | Human GBM patient-derived neurospheres | EGFRvIII amplification | PTEN loss, CDKN2A loss | GBM | 51 | Male | Primary | XRT | Nathanson et al., 2014 |
| HK301 | Human GBM patient-derived neurospheres | EGFRvIII amplification | CDKN2A loss | GBM | 65 | Male | Primary | untreated | Laks et al., 2016 |
| HK296 | Human GBM patient-derived neurospheres | EGFRvIII amplification | PTEN loss, CDKN2A loss | GBM | 76 | Male | Recurrent | XRT, TMZ, Avastin | Laks et al., 2016 |
| HK359 | Human GBM patient-derived neurospheres | EGFRvIII amplification | CDKN2A loss | GBM | 27 | Male | Recurrent | TMZ, XRT, CCNU, CPT-11, carboplatin, avastin | Laks et al., 2016 |
| HK336 | Human GBM patient-derived neurospheres | EGFR wild-type-amplification or gain | PTEN loss, CDKN2A loss | GBM | 65 | Male | Recurrent | TMZ, XRT, Avastin, carboplatin, CCNU | Laks et al., 2016 |
| HK217 | Human GBM patient-derived neurospheres | EGFR wild-type-amplification or gain | PTEN loss, CDKN2A loss | GBM | 81 | Male | Primary | untreated | Laks et al., 2016 |
| HK390 | Human GBM patient-derived neurospheres | EGFR wild-type-amplification or gain | CDKN2A loss | GBM | 73 | Male | Primary | untreated | Laks et al., 2016 |
| HK385 | Human GBM patient-derived neurospheres | EGFR wild-type-amplification or gain | PTEN loss, CDKN2A loss | GBM | 48 | Male | Primary | untreated | Laks et al., 2016 |
| HK254 | Human GBM patient-derived neurospheres | EGFR wild-type-amplification or gain | PTEN loss, CDKN2A loss | GBM | 51 | Female | Recurrent | TMZ, XRT, Avastin | Laks et al., 2016 |
| HK350 | Human GBM patient-derived neurospheres | non-EGFR amplified | PTEN loss, CDKN2A loss | GBM | 81 | Female | Recurrent | TMZ, XRT, Avastin, lapatinib | Laks et al., 2016 |
| HK250 | Human GBM patient-derived neurospheres | non-EGFR amplified | TP53 loss, CDKN2A loss | GBM | 30 | Female | Recurrent | TMZ, XRT | Laks et al., 2016 |
| HK229 | Human GBM patient-derived neurospheres | non-EGFR amplified | PTEN loss, CDKN2A loss | GBM/ Gliosarcoma | 54 | Male | Recurrent | TMZ, XRT | Laks et al., 2016 |
| TS576 | Human GBM patient-derived neurospheres | PDGFRA amplification | PDGFRA amplification | GBM | n/a | n/a | Primary | n/a | Hu et al., 2016 |
| CA718 | Human GBM patient-derived neurospheres | PDGFRA amplification | PDGFRA amplification | GBM | n/a | Female | Primary | nilotinib | Turner et al., 2017 |
| DIPG25 | Human Diffuse Intrinsic Pontine Glioma | H3.3 K27M | TP53 frameshift, MYC amplified | DIPG | 4 | Female | Primary | XRT | Nagaraja et al., 2019 |
| DIPG38 | Human Diffuse Intrinsic Pontine Glioma | H3.1 K27M | TP53 loss, PTEN loss | DIPG | 4 | Female | Primary | XRT, Bevacizumab, Univ. of Pittsburgh vaccine, vorinistat trametinib, pablociclib, everolimus | Nagaraja et al., 2019 |
| DIPG36 | Human Diffuse Intrinsic Pontine Glioma | H3.1 K27M | ACVR1 missense, BCOR frameshift | DIPG | 3 | Female | Primary | XRT, Bevacizumab, Abemaciclib | Nagaraja et al., 2019 |
| DIPG50 | Human Diffuse Intrinsic Pontine Glioma | H3.1 K27M | TP53 loss, RB1 loss | DIPG | n/a | Female | Primary | n/a | Buczkowicz et al., 2014 |

**Document S1.** Evaluation of Fluoxetine in Overall Survival of GBM Patients in Electronic Medical Records from The IBM Marketscan Dataset.

### 1. Data Overview

We used IBM Health MarketScan database[1] (time period between 2003 and 2017), representing time-stamped health insurance claim records of over a half of the US population. The database is a compilation of data provided by over 350 private health insurance providers and from Medicare-eligible individuals. The database includes limited patient-level demographic information (age, sex, geographic area) coupled with patient-specific diagnostics, procedures, laboratory tests, and prescription drugs.

The 2003-2017 version of the database used in this study included 182 million individuals with 51.3% female and 48.7% male enrollees. The mean age at the time of enrollment was 33.6 years (SD: 20 years) and a mean follow up time per enrollee of 2.65 years (SD 2.7 years). Subjects were followed asynchronously and the follow up time varies from few months to several years. The database comes with person-level enrollment information which provides precise start and end dates of the follow up. Some subjects had gaps in their enrollment, *i.e.* their appearance in database spans over two or more disjoint enrollment periods. However, each entry and exit from the database is documented and time-stamped.

Patient-level diagnostic histories are encoded with International Classification of Disease (ICD) (version 9: ICD-9, before October 2015, and version 10: ICD-10, after October 1, 2015). The diagnostic portion of the database contains over 7.6 billion ICD codes. Similarly, procedures performed on patients are represented by corresponding procedure codes (ICD clinical modification codes, Current Procedural Terminology (CPT) codes, and Healthcare Common Procedure Coding System (HCPCS) codes)[2]. The procedures database represents over 8 billion patient-specific events. Time-stamped prescription medications filled by individual patients are represented by their corresponding National Drug Codes (NDCs). Medication part of the database describes over 4.3 billion filled prescription medicines.

### 2. Defining Malignant Neoplasm of Brain

We used ICD-9 codes 191.x and ICD-10 codes C71.x to identify subjects with brain cancer diagnosis. These ICD codes provide location of the cancer within the brain regions but makes no distinction between various forms of brain cancers. Therefore, other proxies such as prescription drugs and performed procedures were used to enrich cohort for glioblastoma multiforme (GBM) patients, as we explain below.

**Sup Table 1** shows the list of ICD codes along with their definition that we used to define the initial study cohort. We identified a total of 165,215 patients with a history of at least one of the diagnostic codes listed in **Sup Table 1**. Briefly, the claims records of patients in this cohort span over 40 million diagnostic codes, over 41 million performed procedures, and over 15 million filled prescription medicines.

| ICD Code | ICD Version | Description |
|---|---|---|
| 191 | ICD-9 | Malignant neoplasm of brain |
| 1910 | ICD-9 | Malignant neoplasm of cerebrum, except lobes and ventricles |
| 1911 | ICD-9 | Malignant neoplasm of frontal lobe |
| 1912 | ICD-9 | Malignant neoplasm of temporal lobe |
| 1913 | ICD-9 | Malignant neoplasm of parietal lobe |
| 1914 | ICD-9 | Malignant neoplasm of occipital lobe |
| 1915 | ICD-9 | Malignant neoplasm of ventricles |
| 1916 | ICD-9 | Malignant neoplasm of cerebellum nos |
| 1917 | ICD-9 | Malignant neoplasm of brain stem |
| 1918 | ICD-9 | Malignant neoplasm of other parts of brain |
| 1919 | ICD-9 | Malignant neoplasm of brain, unspecified |
| C71 | ICD-10 | Malignant neoplasm of brain |
| C710 | ICD-10 | Malignant neoplasm of cerebrum, except lobes and ventricles |
| C711 | ICD-10 | Malignant neoplasm of frontal lobe |
| C712 | ICD-10 | Malignant neoplasm of temporal lobe |
| C713 | ICD-10 | Malignant neoplasm of parietal lobe |
| C714 | ICD-10 | Malignant neoplasm of occipital lobe |
| C715 | ICD-10 | Malignant neoplasm of cerebral ventricle |
| C716 | ICD-10 | Malignant neoplasm of cerebellum |
| C717 | ICD-10 | Malignant neoplasm of brain stem |
| C718 | ICD-10 | Malignant neoplasm of overlapping sites of brain |
| C719 | ICD-10 | Malignant neoplasm of brain, unspecified |

**Supplementary Table 1**. List of ICD-9 and ICD-10 codes used to define brain cancer cohort.

### 3. Workflow for Initial Cohort Selection

In our analysis, GBM patient survival time before all-cause death is the main outcome of interest. However, the information on death status is not available for most enrollees represented in the IBM Health MarketScan database. The only condition in which death of an enrollee is reliably recorded in the database is when patient dies during an inpatient hospital treatment.

We scanned diagnostic histories of all unique enrollees in the MarketScan dataset and identified 378,685 patients who had discharge status “died” upon discharge from hospital. For patients with brain cancer diagnosis, a total of 8,419 patients died in a hospital (**Sup Figure 1**). For the deceased patients, we assumed date of discharge from the hospital as the death date. We restricted our survival time analysis to these 8,419 patients, with an explicit understanding that these patients were likely the sickest ones within our brain cancer cohort. To minimize biases in the analysis, we restricted our brain cancer cohort only to patients with confirmed inpatient death. The known-death-status cohort (*N*=8,419) was described with 2.6 million diagnostics codes, over 2.9 million performed procedures, and nearly 0.7 million filled prescription drugs.

We used other proxies such as prescription drugs and performed procedures to enrich cohort for GBM patients. We scanned diagnostic, procedure, and medication claims of all brain cancer patients to ascertain the timeline of events related to diagnosis and treatments during the follow-up period.

Treatment with temozolomide and radiation therapy and surgical resection of brain tumor is considered a standard of care for GBM patients. We tried to enrich our brain cancer cohort that match these treatment dynamics.

- To enrich for GBM among brain cancer patients, we used NDCs to query patient claims history for medications most frequently used for GBM treatment with or without depression symptoms: temozolomide (217 NDC codes), chemotherapy (2,110 NDCs), fluoxetine (910 NDCs), citalopram (692 NDCs), and escitalopram (338 NDCs, all medication identified from IBM Health MarketScan Redbook 2018 database, see Supplementary Table 1 for details). In addition, we used CPT codes J8700 and J9328 representing oral administration of temozolomide.

- To ascertain radiation therapy, we used CPT code range 77261- 77799, and radiation therapy services HCPCS codes in range G6001 to G6017.

- For detecting surgical resection of brain tumor, we used CPT codes relevant to craniectomy or craniotomy procedures for surgical resection of brain tumor (codes 61510, 61516, 61518, 61520, 61521, 61524, 61526, 61530, 61534, 61536, 61544, and 61545).

- For detecting chemotherapy treatment, we used CPT codes representing “chemotherapy administration and other highly complex drug or highly complex biologic agent administration” in range 96401- 96549; chemotherapy drugs HCPCS code range J9000-J9999, and codes for chemotherapy administration with intravenous infusion and other than infusion technique, G0498, Q0083, Q0084, and Q0085.

- To further detect antineoplastic therapies (radiation therapy, chemotherapy, immunotherapy, or their combinations), we used ICD-9 codes: V58.0, V58.1, V58.11, V58.12, V66.1, V66.2, V67.1, V67.2, and ICD10 codes: Z51.0, Z51.1, Z51.11, Z51.12, and, Z08. These codes were helpful in accurately identifying patients with ongoing or history of cancer or cancer treatments.

- For identifying brain cancer diagnoses, we used ICD-9 codes 191.x and ICD-10 codes C71.x (see **Sup Table 1**). For identifying all other cancer diagnoses, we used entire ICD sub-classification of malignant neoplasms (ICD-9 range 140-209 and ICD-10 code range C00-C96) excluding codes for malignant neoplasm of brain.

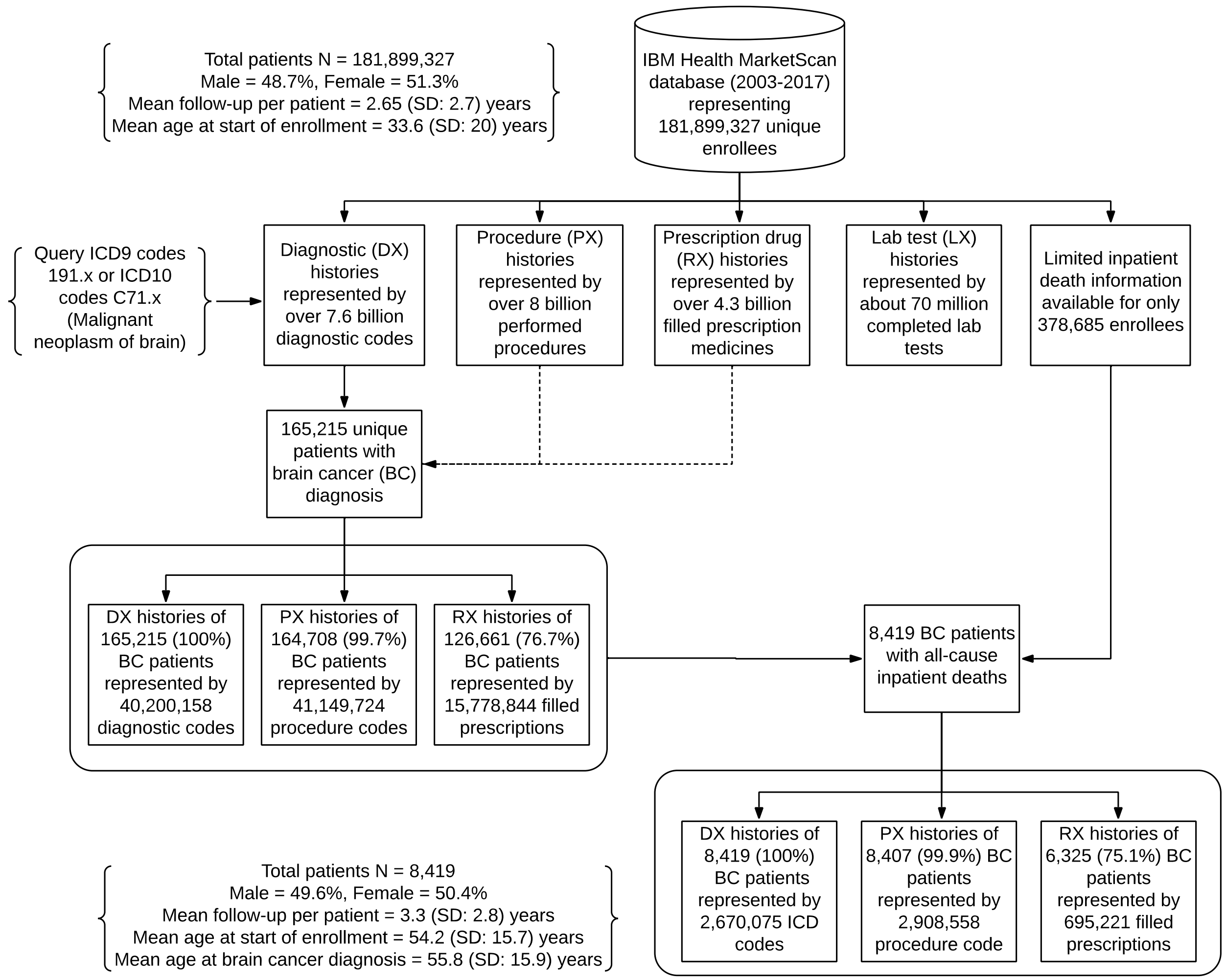


**Supplementary Figure 1.** Illustration of data extraction pipeline and related statistics. The initial target population was patients with at least one claim resulting in brain cancer diagnosis (*N*=165,215). For this subset of population, complete diagnostic, procedures, and prescription histories were obtained (as shown by the dotted lines). Information on patient death was obtained from hospital discharge summaries.

### 4. Exclusion Criteria and Enrichment for Glioblastoma Multiforme

We performed three major pre-processing steps to select the final GBM-enriched cohort.

At the first step, we excluded from the initial brain cancer cohort 4,227 patients who had incomplete, missing, or irrelevant health histories, see **Sup Figure 2**. We excluded from the cohort all enrollees who were under 18 years of age at the time of diagnosis, focusing on older enrollees as target population. We then excluded data for all patients who had less than one year of enrollment documented, see **Sup Figure 2**, to make sure that health histories were long enough for a meaningful downstream inference. (This is necessary because some people switch or cancel insurance at various points in time, creating gaps in their documented health histories.) For the same reason we excluded from the cohort all patients with gaps in their documented health histories spanning more than a month.

In the IBM Health MarketScan dataset, at any given time, about 75% of all enrollees had prescription drugs benefit with their insurance coverage. Because we used chemotherapy drugs as cancer diagnosis proxies, and antidepressant drugs as exposures, it was critical for us to retain only those patients had full medication coverage benefit during their entire enrollment period. Therefore, we excluded 1,365 patients who did not have medication coverage benefits for their entire follow up period.

In the second pre-processing step, we excluded 3,073 patients who had either history of or ongoing cancer treatment, or have any cancer that could be metastatic, before the index brain cancer diagnosis. The index brain cancer diagnosis here refers to the appearance of first brain cancer ICD code in the patient diagnostic histories. We also excluded patients who did not have at least three months of history prior to index brain cancer diagnosis. This was done to filter out patients who may join insurance (become visible in our database) with some ongoing treatment and diagnoses that we cannot ascertain.

ICD-9 and ICD-10 codes do not differentiate between various types of malignant brain neoplasms. Instead, they provide information on the affected brain regions, thus somewhat limiting our ability to differentiate GBM from other tumors. Also, ICD codes make no differentiation between initial tumor diagnosis or a relapse. Given these limitations, we made an effort to enrich our cohort for GBM using medication and procedure proxies. Specifically, we used records of temozolomide (TMZ) medication, of radiation therapy (RT), and of surgical resection of tumor as GBM proxies to enrich the cohort for GBM patients. We removed 881 patients who did not meet the GBM enrichment criteria (**Sup Figure 2**).

The final GBM-enriched cohort included 238 patients all of which died in hospital after receving standard GBM care. Out of these 238 patients, 56 patients were treated with SSRI antidepressants, in addition to treatments included in the standard of care for GBM. Out of these 56 patients, 10 were treated with fluoxetine (8 with fluoxetine only, 1 with fluoxetine and escitalopram, and 1 with both fluoxetine and citalopram), 13 with citalopram (9 citalopram only, and 3 with citalopram and escitalopram), and 38 with escitalopram (34 with escitalopram only). Note, this cohort was selected to represent exposures to antidepressants *after* the index GBM diagnosis. The question we asked was: do (any of) antidepressants change survival odds when prescribed in addition to standard care of GBM? We did not consider exposure to antidepressants prior to GBM diagnosis.

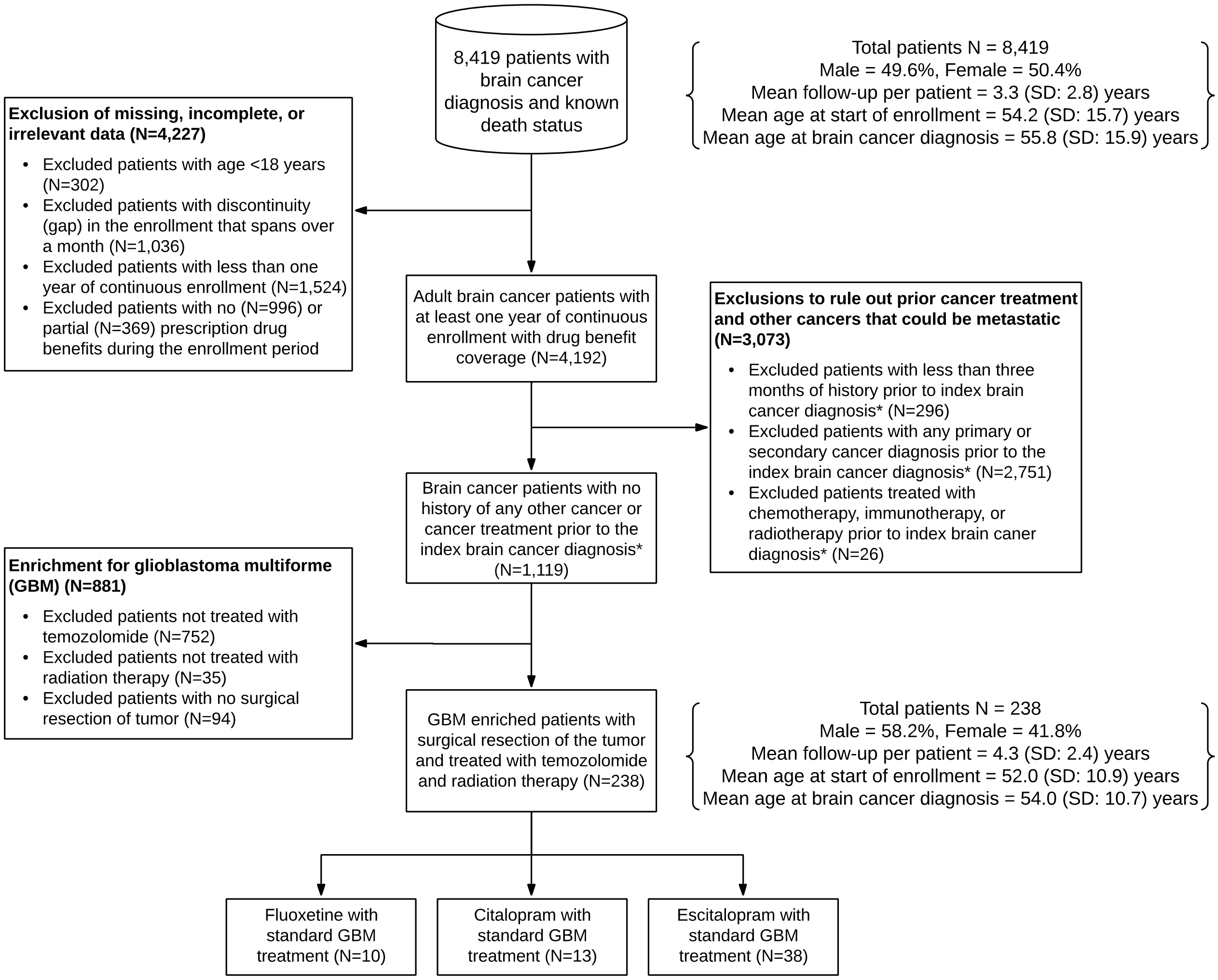


** Index brain cancer diagnosis refers to the appearance of first brain cancer ICD code in the IBM Health MarketScan diagnostic claims records of the patient.*

**Supplementary Figure 2.** Illustration of inclusion and exclusion criteria to refine the brain cancer cohort and enrichment for glioblastoma multiforme (GBM). These inclusion and exclusion steps were performed to minimize potential biases that are usually inherent to the observational dataset.

### 5. Statistical Analysis

We performed a version of *time to event* analysis by comparing survival probabilities between exposed and unexposed groups. The event of interest was all-cause inpatient death following GBM diagnosis. We used the Kaplan-Meier (KM) method[3] to estimate the survival function for GBM patients treated with one of the three SSRI antidepressants against those with standard GBM treatment but no SSRI antidepressant exposure. Further, extended Cox proportional hazard regression model with time-dependent variable[4,5] was used to assess the variation in hazards associated within three SSRI antidepressants exposure. Separate experiments for each SSRI antidepressants was conducted to estimate the hazards ratio (HR) of all-cause death among the GBM patients.

As explained in the previous section, the described GBM-enriched cohort included 238 patients, all having brain cancer diagnosis followed by surgical resection of tumor and treated with temozolomide and radiation therapy. For each GBM subject, we computed the number of days from index GBM diagnosis to the day the patient died in a hospital. To adjust results for the *immortal time bias*[6,7], SSRI antidepressants exposure was computed as a time interval-dependent value[8] – the patient was considered unexposed to antidepressant until the first dispensed prescription of corresponding SSRI, and considered exposed only after medication was being prescribed. The start date of the SSRI antidepressant exposure was the date of first SSRI antidepressant prescription dispensed after index GBM diagnosis.

For proper comparison, we used consistent controls for each of the three SSRI antidepressant exposure models. The control group included 182 patients that underwent standard GBM treatment but were not exposed to any of the three SSRI antidepressants considered in this study.

We compared the survival curves for the two groups (SSRI antidepressant exposure, no SSRI antidepressant exposure) and also compared them using time dependent Cox proportional hazards regression and adjusting for confounders such as age and sex.

We implemented Cox PH regression in R, using "survival" package[9] and "coxph" function with time dependent exposure modelling[10]. Multiple observations from the same patient (one prior to SSRI antidepressant exposure and one after SSRI antidepressant exposure) were indicated by the patient-level identifier in the mode to compute a robust (cluster) variance for the model. The validity of proportional hazards assumption was evaluated using Schoenfeld test[11,12] performed on individual predictors and the over model.

### 6. Results

A visual inspection of time-dependent KM curves suggests that patients treated with fluoxetine have survival advantage over those not treated with any of the three SSRI antidepressants considered in this study (**Figures 6A-6D**). Those treated with fluoxetine survived longer (median survival 545 days) compared to those not treated with fluoxetine (median survival 318 days). The age and sex adjusted HR of all-cause death in fluoxetine treated group was 0.42 [95% CI, 0.20 – 0.88], p = 0.02 compared to the control group (**Sup Figure 3**). There was no significant difference in all-cause hazards of death in citalopram or escitalopram vs. the control group (**Sup Figures 4** and **5**). There was no significant difference in HR for sex but the HR of all-cause death increases by 3% with increasing age (**Sup Figures 3**, **4**, and **5**).

We validated the proportional hazards assumption with Schoenfeld test for all three models and found no evidence of violation of the PH assumption (**Sup Figures 3**, **4**, and **5**).

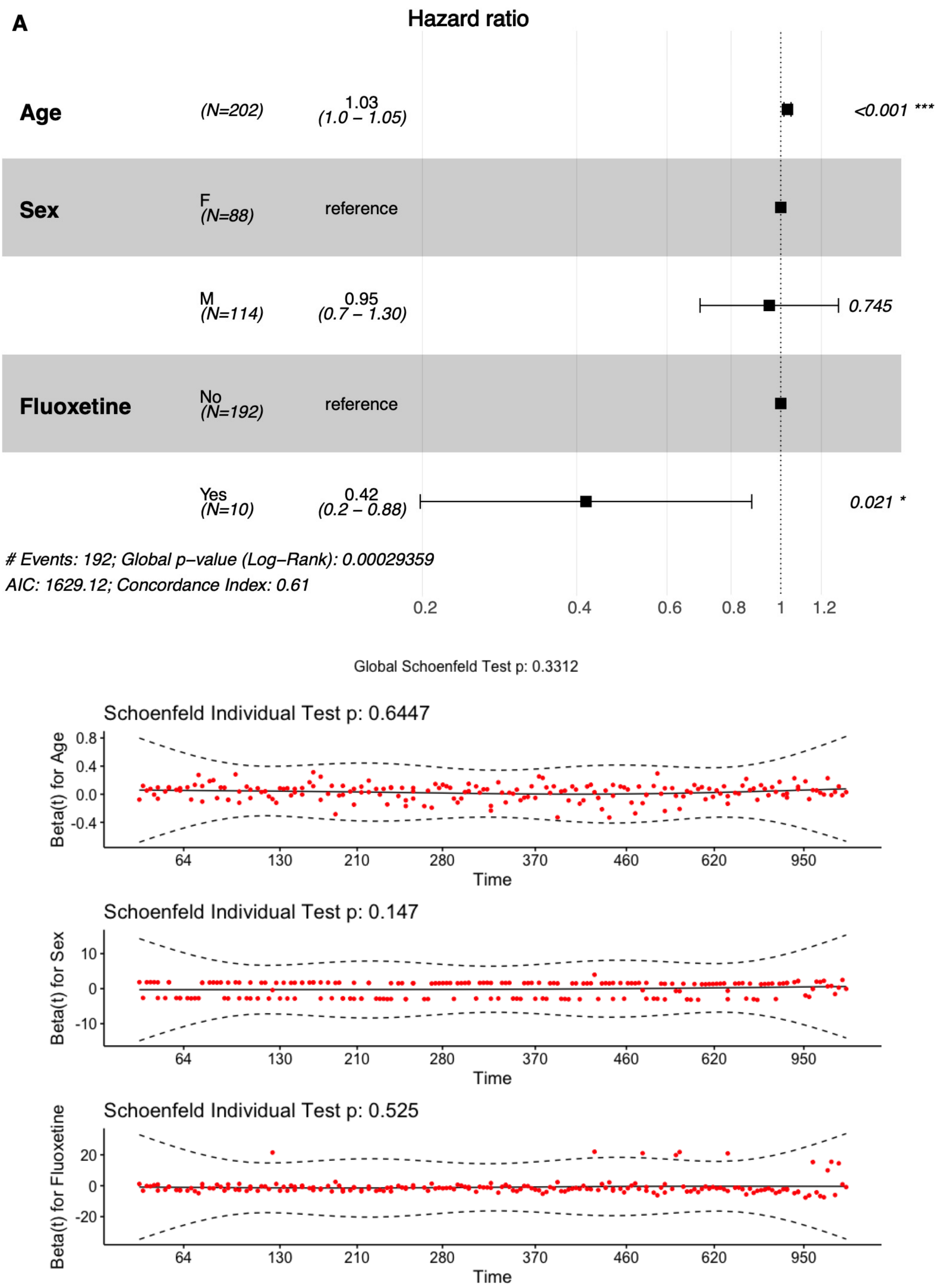


**Supplementary Figure 3.** Top: Estimated hazard ratios (using Cox proportional hazards regression) for all-cause death in GBM-enriched cohort. Fluoxetine exposure was treated as time-dependent variable. Bottom: Validity of the proportional hazards assumption was evaluated with Schoenfeld global and individual tests (the null hypothesis corresponding to the assumption is not rejected).

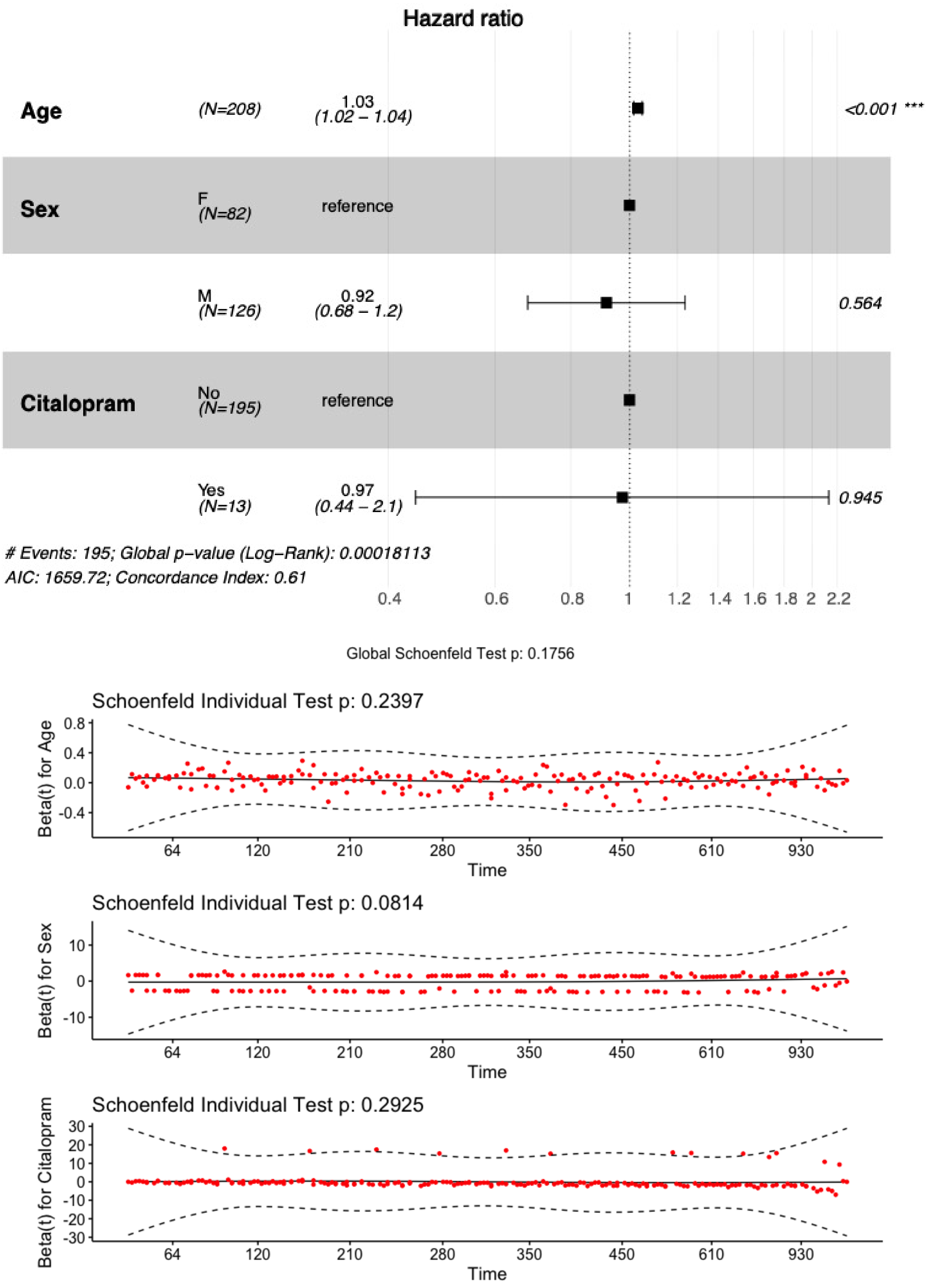


**Supplementary Figure 4.** Top: Estimated hazard ratios (using Cox proportional hazards regression) for all cause death in GBM enriched patients. Citalopram exposure was treated as time-dependent variable. Bottom: Validity of the proportional hazards assumption was evaluated with Schoenfeld global and individual tests.

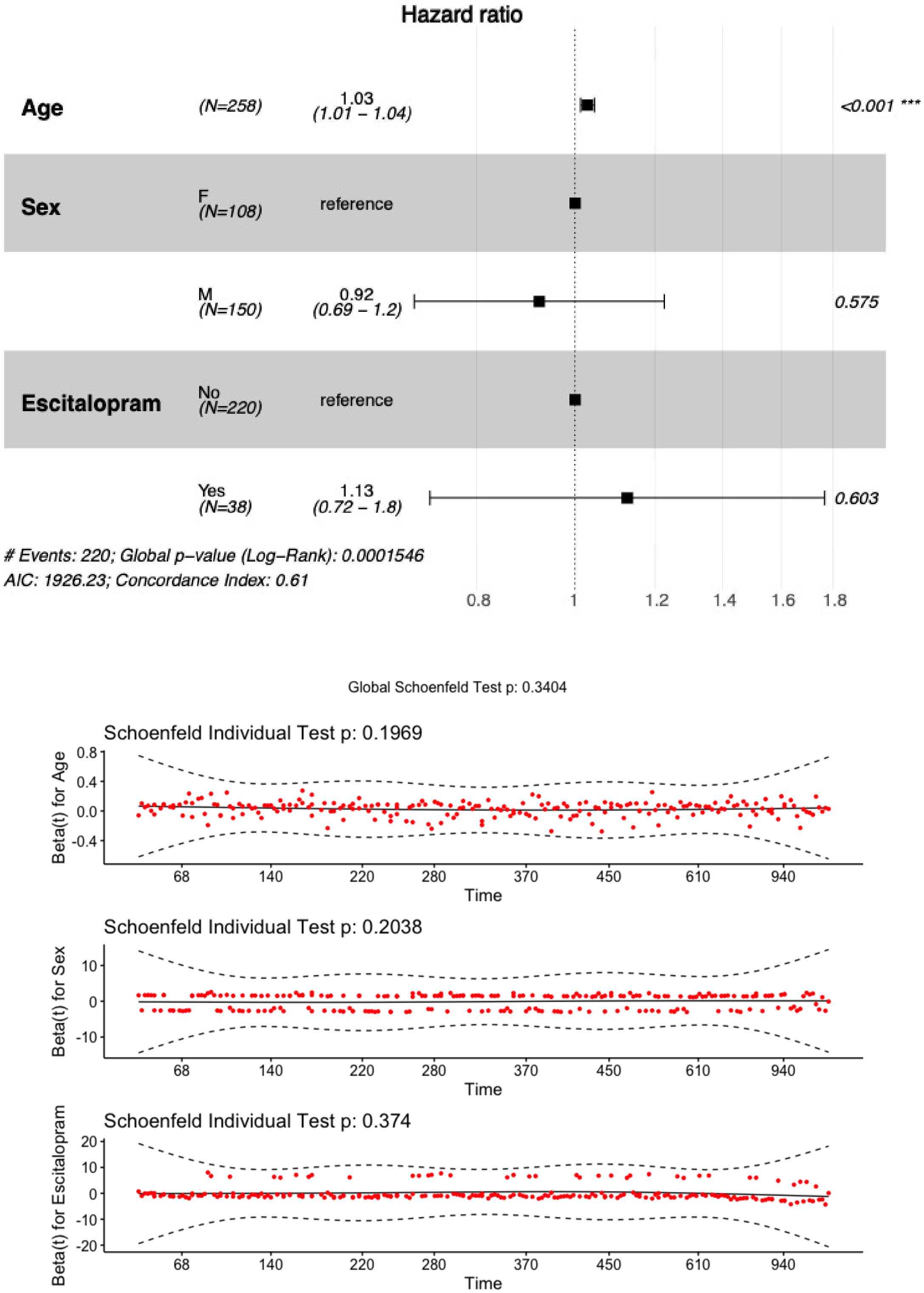


**Supplementary Figure 5.** Top: Estimated hazard ratios (using Cox proportional hazards regression) for all cause death in GBM enriched patients. Escitalopram exposure was treated as time-dependent variable. Bottom: Validity of the proportional hazards assumption was evaluated with the Schoenfeld global and individual tests.
